# The axonal ER couples translation and secretion machineries for local delivery of axonal transmembrane proteins to promote axonal development

**DOI:** 10.1101/2025.09.09.674816

**Authors:** Ha H. Nguyen, Noortje Kersten, Chun Hei Li, Huey de Jong, Tanuj Arora, Themistoklis Liolios, Dan T. M. Nguyen, Maarten P. Bebelman, Maarten Altelaar, Max Koppers, Ginny G. Farías

## Abstract

Neuronal development and function rely on the polarized sorting of transmembrane proteins (TMPs). Most TMPs follow a conventional secretory pathway confined to the cell soma, where ER-translated TMPs exit the ER through ERES to reach the Golgi, prior to PM delivery. We recently found that the axonal ER regulates local translation; however, how this is linked to secretion remains unknown. Here, we find that axonally translated TMPs exit the ER and reach the axonal PM via an ERES-dependent/Golgi-independent route. Intriguingly, we uncover a feedback loop linking translation and secretion at axonal ERES. HDLBP-dependent TMP synthesis regulates ERES formation, and axonal ERES in turn modulate TMP translation. Subsequently, the NRZ-SEC22B tethering complex links local TMP ER exit to PM delivery, dependent on ER-PM contacts. Axonal ERES and these associated molecular players support axon growth and bouton assembly. We propose a novel mechanism in which coupling of axonal TMP synthesis and secretion is essential for neuronal development.

## Introduction

Neurons are highly polarized cells comprising a soma, multiple dendrites and a long axon. These different neuronal compartments exhibit distinct molecular components and functions, which are important for neurons as signal integration centers. In humans, axons can extend up to one meter in length, resulting in a total surface area that may be thousands of times larger than that of the soma. This large surface area demands substantial amounts of membranes and proteins in the axon^1,2^. Additionally, during development and synaptic activity, axons need to respond quickly to intracellular signals and ongoing changes in their external environment to ensure growth and plasticity^3-5^. Therefore, precise spatiotemporal regulation of the axonal proteome is required.

RNA localization and local translation allow axons to regulate their proteome in response to local demand, playing a vital role during axonal development and synaptic activity^6^. Axons possess the necessary translational machinery^7-9^ and studies employing compartment-specific transcriptome and translatome analyses have identified thousands of mRNAs localized and translated within them^10,8,11^. Interestingly, a significant portion of these are mRNAs encoding transmembrane proteins (TMPs)^12,13,8,14^. These include neuronal cell adhesion molecules, guidance and signaling receptors, and synaptic proteins. This suggests that translation of TMPs occurs along the axon and has an impact during different stages of neuronal development. Most TMPs follow a conventional secretory pathway, where they are translated at the rough endoplasmic reticulum (ER), exit through ER exit sites (ERES) and are sorted to the Golgi before being delivered to their target locations at the plasma membrane (PM). However, major components of the secretory pathway, including the rough ER and the Golgi are restricted to the somatodendritic domain in CNS neurons^15,16^, raising fundamental questions about local TMP biogenesis and sorting in axons. Notably, axons harbor extensive ER tubules that associate with ribosomes to regulate local translation in an extrinsic cue-dependent manner^17^. Many mRNAs encoding TMPs are associated with P180/RRBP1, an axonally-enriched ER protein that facilitates the interaction of ribosomes with the axonal ER to regulate local translation^17^. Yet, how axonally synthesized TMPs exit the ER and reach the PM, and how local translation and secretion are linked to ensure proper local delivery of neosynthesized TMPs to the PM remain elusive.

In this study, we show mRNA localization and local translation of different TMPs along the axon shaft and at branching points and axon tips. These neosynthesized TMPs can exit the axonal ER and reach the PM, independently of the Golgi/Golgi-derived organelles, but dependent on axonal ERES. Interestingly, we uncover that the ER-translation regulator HDLBP and the NRZ-SEC22B tethering complex are proximal to axonal ERES, locally linking translation and secretion to the PM at axonal ERES. Mechanistically, we found a feedback loop where HDLBP controls local TMP translation and ERES formation, while axonal ERES regulate local TMP translation. Consecutively, SEC22B, involved in TMP ER exit and in ER-PM contact sites, together with the NRZ complex link ER exit of TMPs with local secretion to the PM. Finally, we show that axonal ERES and identified players contribute to axonal growth and bouton assembly. Together, we propose a novel mechanism coupling axonal translation and secretion of TMPs to the PM, which is critical for axon development.

## Results

### Local TMP translation occurs along the axonal shaft and at axonal branches and tips

Many mRNAs encoding TMPs have been identified in various axonal transcriptome and translatome datasets^8,11,12,14^. Here, we set out to visualize axonal mRNA localization and/or local translation of the synaptic protein SYT1 and the adhesion molecules NRXN1α and L1CAM in rat hippocampal neurons at day-in-vitro (DIV) 10. These proteins play critical roles in axonal development and maintenance^18-20^. To visualize TMP mRNAs, we used the PP7-PCP system, introducing PP7 RNA stem-loops into the 3’UTR of SYT1 and L1CAM mRNAs and co-expressing them together with a 3x-HaloTag fused to the PP7 coat protein for their detection (**Fig. 1a**)^21^. mRNA puncta for SYT1 and L1CAM were observed along the axon (**Fig. 1b**). To visualize translation of TMPs in the axon, we performed puromycin labeling coupled with a proximity ligation assay (puro-PLA) (**Fig. 1c**). In brief, we used two primary antibodies, one recognizing puromycin and the other recognizing the TMP protein of interest (POI). When both antibodies bind the same nascent TMP, oligonucleotide-coupled secondary probes hybridize to form a DNA bridge that supports rolling-circle amplification, yielding fluorescent puncta that report local translation sites of the POI^22-24^. We visualized the axonal translation of SYT1, NRXN1α and L1CAM. Although these TMPs are actively produced in the cell soma, we found that their local translation can also be observed in the axon (**Fig. 1d, f**). The number of Puro-PLA puncta along the axon was normalized to total axon area using the axonal neurofilament marker SMI-312 (**Fig. 1d-g**). There was a significant difference in the number of puncta between neurons treated with puromycin for 10 minutes compared to those pre-treated with the translation inhibitor anisomycin prior to puromycin incubation, indicating the specificity of this assay (**Fig. 1d-g**). Since external cues have been shown to increase local protein synthesis and neuronal activity^25,26,9,27^, we stimulated neurons with BDNF for 30 minutes before puromycin labeling. Interestingly, upon BDNF stimulation, there was a significant increase in local SYT1 translation (from 2.679 to 3.407 puncta per 100 μm^2^), indicating translation of SYT1 can be cue-induced (**Fig. 1d, e**). On the other hand, BDNF stimulation caused only a slight increase in the number of nascent NRXN1α protein (**Fig. 1f, g**). We also observed SYT1 being synthesized at branching points and axon tips (**Fig. 1h**), consistent with ER-ribosome interactions enriched in these axonal sub-compartments^28,29,17^. Furthermore, axonal TMP synthesis was not limited to rat hippocampal neurons, as local translation of SYT1 (**Fig. 1i**), NRXN1α (**Fig. 1k**) and L1CAM (**Fig. 1m**) was also detected in axons of human cortical iNeurons at DIV24, with abundances comparable to those in primary rat neurons (**Fig. 1j, l, n**). These results show that different TMPs can be locally synthesized along the axonal shaft and at axonal branches and tips, and that their translation increases upon neuronal stimulation.

**Fig. 1:**
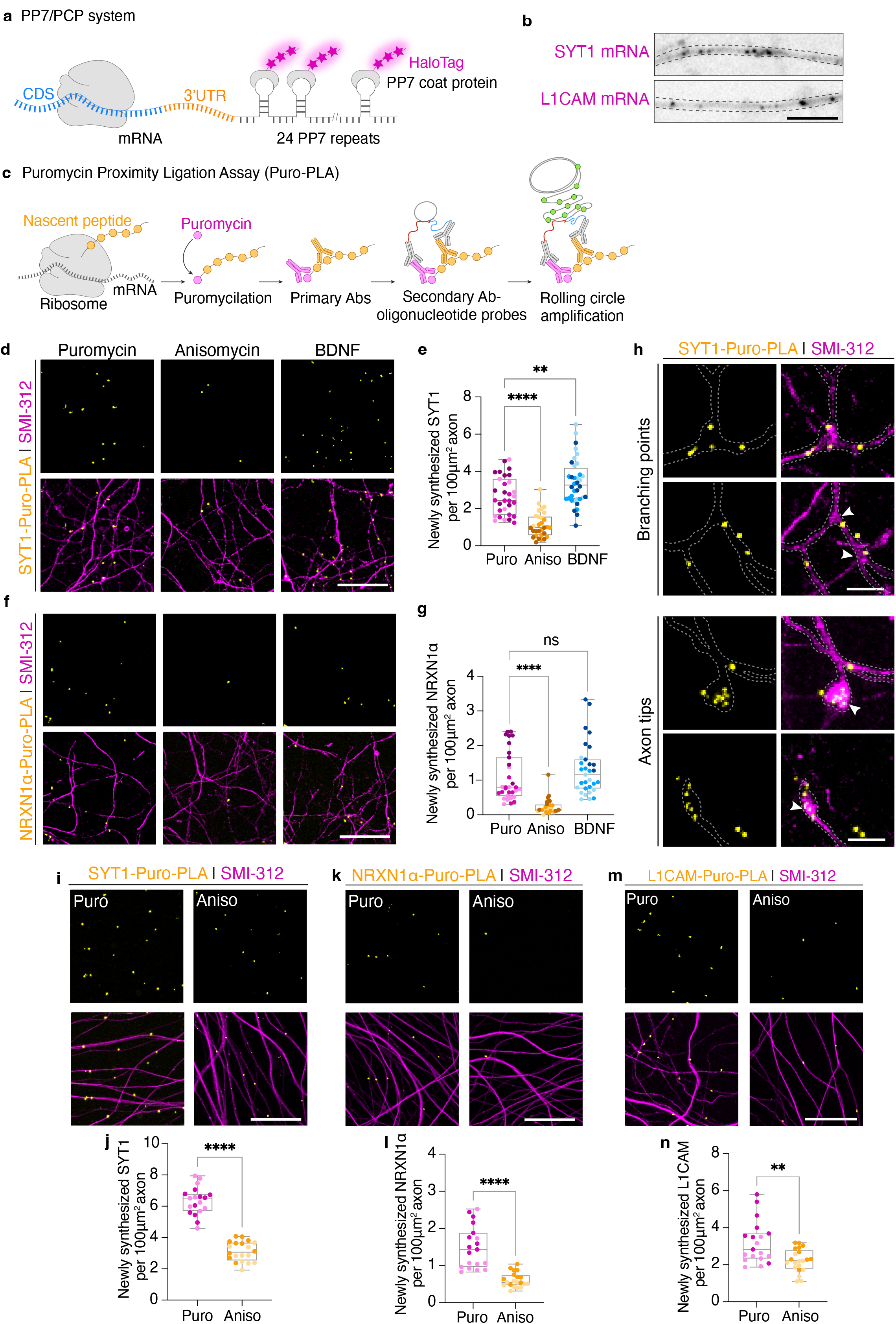
Axonal TMPs are locally synthesized in the axon of primary rat and iPSC-derived human CNS neurons. **a**, Schematic representation of the PP7/PCP system consisting of 24 RNA stem loops inserted in the 3’UTR of the mRNA of interest and HaloTag-PP7 coat protein for its detection. **b**, Representative images of SYT1 and L1CAM mRNA in the axon of primary rat hippocampal neurons at DIV10. **c**, Schematic representation of the puro-PLA assay. **d-g**, Representative images of SYT1 (**d**) and NRXN1α (**f**) translation sites in DIV10 primary rat hippocampal axons labelled with SMI-312, under only puromycin labeling, anisomycin pretreated, or BDNF pretreated conditions. Quantification of newly synthesized SYT1 (**e**) and NRXN1α (**g**) per 100 μm^2^ axon. Data were collected from 3 independent experiments (N=3). **h**, Representative images of SYT1 translation sites in branching points or axon tips, indicated by white arrow heads. **i-n**, Representative images of SYT1 (**i**), NRXN1α (**k**), and L1CAM (**m**) translation sites in DIV24 iPSC-derived human cortical neurons under only puromycin labeling or anisomycin pretreated conditions. Quantification of SYT1 (**j**), NRXN1α (**l**) and L1CAM (**n**) translation sites per 100 μm^2^ axon, (N=2). Data are presented as box-and-whisker plots in (**e, g, j, l, n**). Individual data points each represent an axon area and each color per condition represents an independent experiment. ns = non-significant, **p < 0.01, ****p<0.0001 comparing conditions with control using ordinary one-way ANOVA test, followed by a Dunnett’s multiple comparisons test (**e**), Kruskal-Wallis test followed by Dunn’s multiple comparisons test (**g**), or unpaired t-test (**j, l, n**). Scale bars represent 20 μm in (**d, f, i, k** and **m**) and 5 μm in (**b, h**).

### Newly synthesized TMPs exit the axonal ER and are delivered to the PM

Since we observed translation of TMPs occurring along the axon, we wondered whether these could be locally secreted. Previous studies have shown a role for Golgi outposts, Golgi satellites, and the ER-Golgi intermediate compartment (ERGIC) in local TMP secretion in dendrites^30-32^. Thus, we first examined different markers for Golgi and Golgi-derived/associated compartments. Although these compartments were observed in the soma and dendrites, they were absent from axons (Supplementary **Fig 1a-d**). To determine whether axonally synthesized TMPs have the capability to be secreted from the axonal ER, we used the Retention Using Selective Hook (RUSH) system. This method allows the retention of POIs in the ER and their synchronous release upon the addition of biotin, to follow their sorting^33^. We first used the ER retention sequence KDEL or the invariant chain of class II MHC as a hook coupled to Streptavidin, and our protein of interest (SYT1, NF186 or L1CAM) coupled to a Streptavidin binding peptide (SBP) and an mNeonGreen (mNG) fluorescent tag for cargo visualization (**Fig. 2a**). We found that at timepoint 0 (no biotin), our TMPs (also referred to as cargoes) are properly retained at the ER, with an enrichment in the somatodendritic domain (**Supplementary Fig. 2a-c**). Upon addition of biotin for 15 minutes, we observed their conventional sorting to the somatic Golgi, followed by their localization to specific compartments after 24h. SYT1 properly reached synaptic vesicles, NF186 localized to the axon initial segment, and L1CAM distributed along the axonal PM and vesicles (**Supplementary Fig. 2a-c**). After 4h release, we observed SYT1 appear as puncta along the axon, with an average of 33 SYT1-positive vesicles per 100 μm axon (**Supplementary Fig. 2d, e**).

**Fig. 2:**
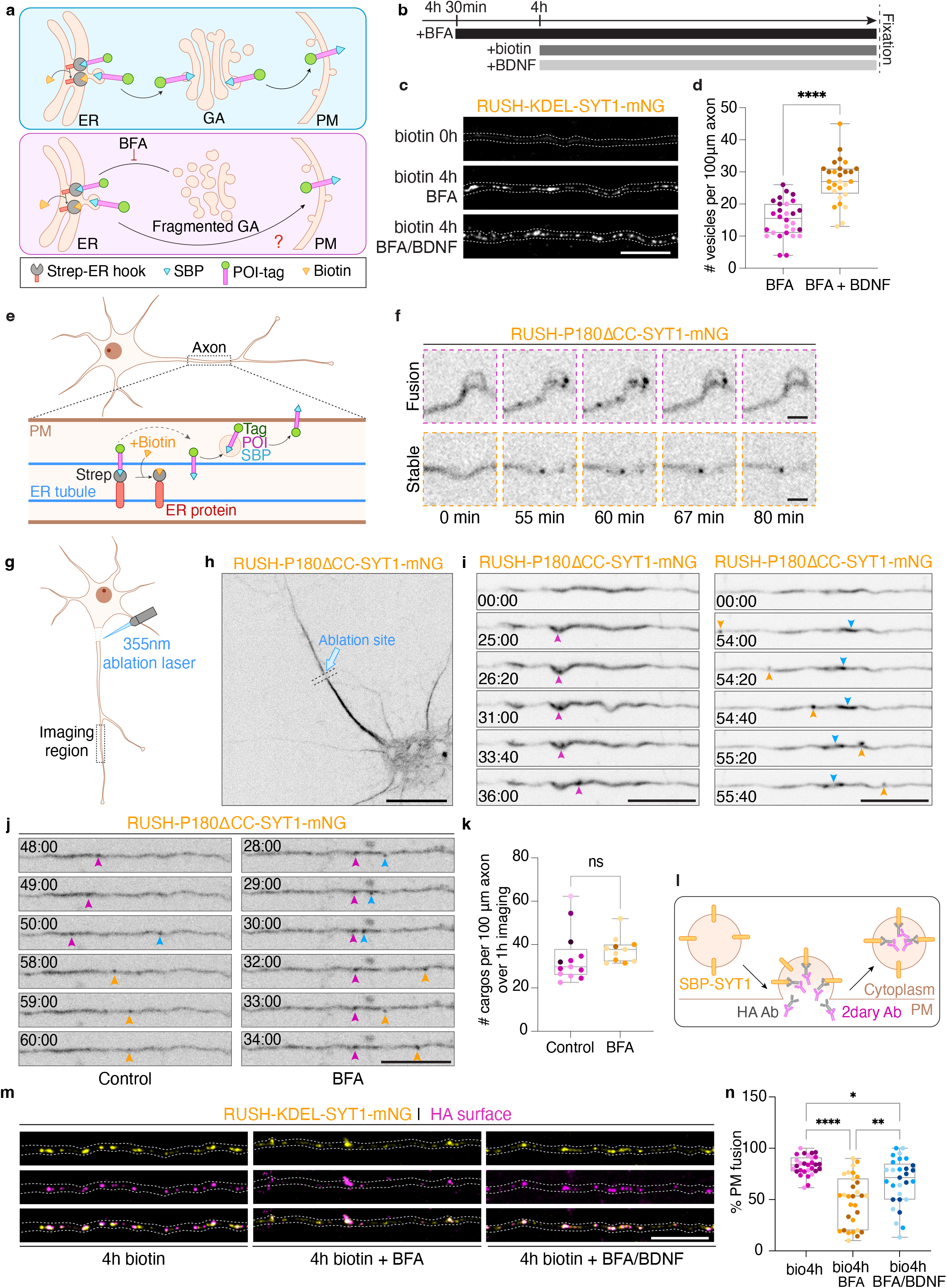
Locally translated TMPs undergo unconventional secretion from the axonal ER to the PM. **a**, Schematic representation of the RUSH system to track cargo trafficking from the ER. Upon biotin addition, cargoes are released, accumulated at the Golgi and sorted to the PM (upper panel). BFA treatment is used to block ER-Golgi trafficking and to assess if cargo can exit the ER and reach the PM (lower panel). **b**, Schematic showing the timeline of different treatments before fixation. **c**, Representative images of RUSH-SYT1 cargo distribution in DIV10-11 axons at timepoint 0, 4h biotin release with BFA, and 4h biotin release with BFA and BDNF. **d**, Quantification of RUSH-SYT1 cargo numbers per 100 μm axon (N=3). **e**, Schematic representation of the RUSH system with axonal ER hook. **f**, Live cell still images showing RUSH-SYT1 exit from the ER in the distal axon at 1-minute intervals for 120 minutes. **g**, Schematic representation of the RUSH system with axonal ER hook in combination with axon severance by photo ablation using 355nm laser. **h**, Representative image of a neuron expressing the RUSH-P180ΔCC-SYT1 system after axon cutting. The blue arrow points to ablation site. **i**, Live cell still images showing two axon segments imaged at 20-second intervals for 60 minutes. Arrowheads point to new cargo exiting the axonal ER. **j**, Live cell still images showing the release of RUSH-SYT1 with P180ΔCC hook in desomatized axons in the absence and presence of BFA. At approximately 15 minutes after axon cutting, images were acquired at 1-minute intervals for 60 minutes. Arrowheads point to new cargoes exiting the axonal ER. **k**, Quantification of the cumulated number of cargos exiting the ER over 1 hour imaging in 100 μm axon (N=5, control, N=4, BFA). **l**, Schematic representation of RUSH-SYT1 cargo release in combination with HA surface labeling. An HA tag is added in the luminal/extracellular side of SYT1. Upon cargo fusing with the PM, HA antibody binds to this epitope and gets endocytosed in synaptic vesicles. **m**, Representative images of DIV10-11 neurons expressing RUSH-SYT1 after 4h biotin release, 4h biotin release and pre-treated with BFA alone, or 4h biotin release together with BFA and BDNF. Live cell surface labeling for HA-tag inserted in the luminal/extracellular domain of RUSH-SYT1, allows for the visualization of SYT1 fusion with the PM. **n**, Quantification of the percentage of released cargoes fusing with the PM (N=3). Data are presented as box-and-whisker plots (**d, k, n**). Individual data points each represent a neuron, and each color per condition represents an independent experiment. *p < 0.05, **p < 0.01, ****p<0.0001 comparing conditions using unpaired t-test (**d**) or ordinary one-way ANOVA test followed by Tukey’s multiple comparisons test (**n**). ns= non-significant, p = 0.0720 comparing total number of cargoes exiting the ER in desomatized axon in control and BFA treated conditions over 1h imaging using Mann-Whitney test (**k**). Scale bars represent 10 μm in **c, h, i, j and m** and 2 μm in **f**. See also Supplementary Figs **1-5** and Supplementary Videos **1-3**.

To study unconventional secretion, independent of somatodendritic ER-Golgi sorting, we used Brefeldin A (BFA). BFA targets Arf1 GTPase, preventing COP-I binding to the Golgi, which eventually leads to overall ER-Golgi trafficking defects and Golgi fragmentation (**Fig. 2a; Supplementary Fig. 3a, b**)^34^. For the RUSH system, BFA was added to the culture medium for 30 minutes, immediately followed by the addition of D-biotin for 4 hours (**Fig. 2b**). We first validated that BFA was sufficient to block ER-Golgi trafficking. The conventional somatodendritic TMP transferrin receptor (TfR)^35,36^ was efficiently retained in the ER upon BFA treatment (**Supplementary Fig. 3c**). We then analyzed axonal TMPs upon BFA treatment. We found that although SYT1 signal was detected in the somatic ER, a portion of SYT1 was still secreted independently of the Golgi, with an average of 15 vesicles per 100 μm axon (**Fig. 2c, d; Supplementary Fig. 3d, e**). Similar results were observed for NF186 and L1CAM (**Supplementary Fig. 3d, e**). Consistent with Golgi-independent secretion, we did not find BFA-sensitive COPI components along the axon (**Supplementary Fig. 3f, g**). Remarkably, neuronal stimulation with BDNF increased the number of Golgi-bypassing cargoes, from ± 15 cargoes to ± 27 cargoes per 100 μm axon (**Fig. 2c, d**). Live cell imaging during BFA incubation showed no clear translocation of Golgi bypassing vesicles from the soma to the axon, suggesting these TMPs are released from the axonal ER (**Supplementary video 1**). To further confirm this, without blocking conventional secretion, we utilized a truncated version of the axon-enriched ER-protein P180 as a hook (Strep-P180ΔCC) (**Fig. 2e; Supplementary Fig. 4a**)^37^. We observed ER bending and cargo budding from the ER in the distal axon after biotin addition (**Fig. 2f**). Some SYT1 vesicles became diffusive, while others underwent short-range movements or remained stably attached to the ER (**Fig. 2f; Supplementary video 2**). To verify that vesicles present along the axon locally exit the axonal ER, we performed axon cutting with a 355nm photoablation laser prior to biotin-induced ER release (**Fig. 2g, h; Supplementary Fig. 4b, c**). Similar to intact axons, we observed local TMP exit from the ER in desomatized axons (**Fig. 2i; Supplementary Video 3**). Importantly, we found a similar number of cargoes exiting the axonal ER over 1h imaging in the absence or presence of BFA in desomatized axons. This indicates that local cargos leave the axonal ER via an unconventional route, without being sorted to any Golgi-related compartment. (**Fig. 2j, k**).

To determine whether these cargoes are inserted at the PM in a similar way as conventional soma-derived cargoes, we performed HA live surface labeling with RUSH-KDEL-SYT1-mNG, which contains an HA tag in the luminal/extracellular domain of SYT1 (**Fig. 2l**). This assay allows us to label SYT1 cargoes once they are delivered to the PM. Of note, Golgi-bypassing SYT1 cargoes reach synaptic vesicle compartments (**Supplementary Fig. 5a, b**), which undergo synaptic vesicle recycling at the PM^38,39^. Therefore, the HA signals are expected to reside predominantly in synaptic vesicles as puncta, instead of being diffusive along the PM. We found that under BFA treatment, newly secreted Golgi-bypassing SYT1 cargoes could still reach the PM, although to a lesser extent than conventional soma-derived cargoes (**Fig. 2m, n**). Interestingly, BDNF stimulation increased Golgi-bypassing cargoes fusing with the PM (**Fig. 2m, n**).

Altogether, these different assays provide evidence that TMPs can locally exit the axonal ER and reach the PM via a Golgi-independent mechanism which can be modulated by neuronal stimulation.

### Axonal ERES are required for local secretion and the ERES proxeome identifies key candidates linking local TMP synthesis and secretion

We next sought to gain insights into how locally synthesized TMPs exit the ER in the absence of Golgi-related compartments. Previous studies have shown that ERES components localize to the axon^40,41^. Consistent with this, we found different ERES components in the axon of rat hippocampal neurons at different stages of development (**Fig. 3a; Supplementary Fig. 6a-d**). Endogenous staining of SEC31A and overexpression of V5-tagged SEC31A showed a similar abundance of axonal ERES puncta (**Supplementary Fig. 6e, f**). We found that ERES components highly colocalized with each other in the axon (**Fig. 3a, b**). To determine if these ERES components are indeed associated with the ER and are not COPII-decorated vesicles, we performed live cell imaging for the ERES marker GFP-SEC23A. Most puncta exhibited little mobility along the axon (**Supplementary Fig. 6g-i**). Importantly, BFA treatment did not interfere with the presence of ERES components in the axon or their dynamics (**Supplementary Fig. 6j-l**). Using a SoRa spinning disk super-resolution microscope, we further observed that ERES remained stably associated with tubular axonal ER, with short-range movements accompanying local tubular ER rearrangement (**Supplementary Fig. 6m; Supplementary Video 4**). BDNF treatment increased number of ERES along the axon, possibly to accommodate enhanced ER exit of newly translated TMPs (**Fig. 3c, d**). To examine whether the axonal ER is required for distribution of ERES along the axon, we removed the ER from the axon. For this, we used an established Strep/SBP heterodimerization system that relocates the axonal ER to the somatodendritic domain via the minus-end directed KIFC1 motor (**Fig. 3e**)^37,42,17^. ER tubule removal from the axon resulted in removal of axonal ERES components, indicating that they are tightly associated to the axonal ER (**Fig. 3f, g**).

**Fig. 3:**
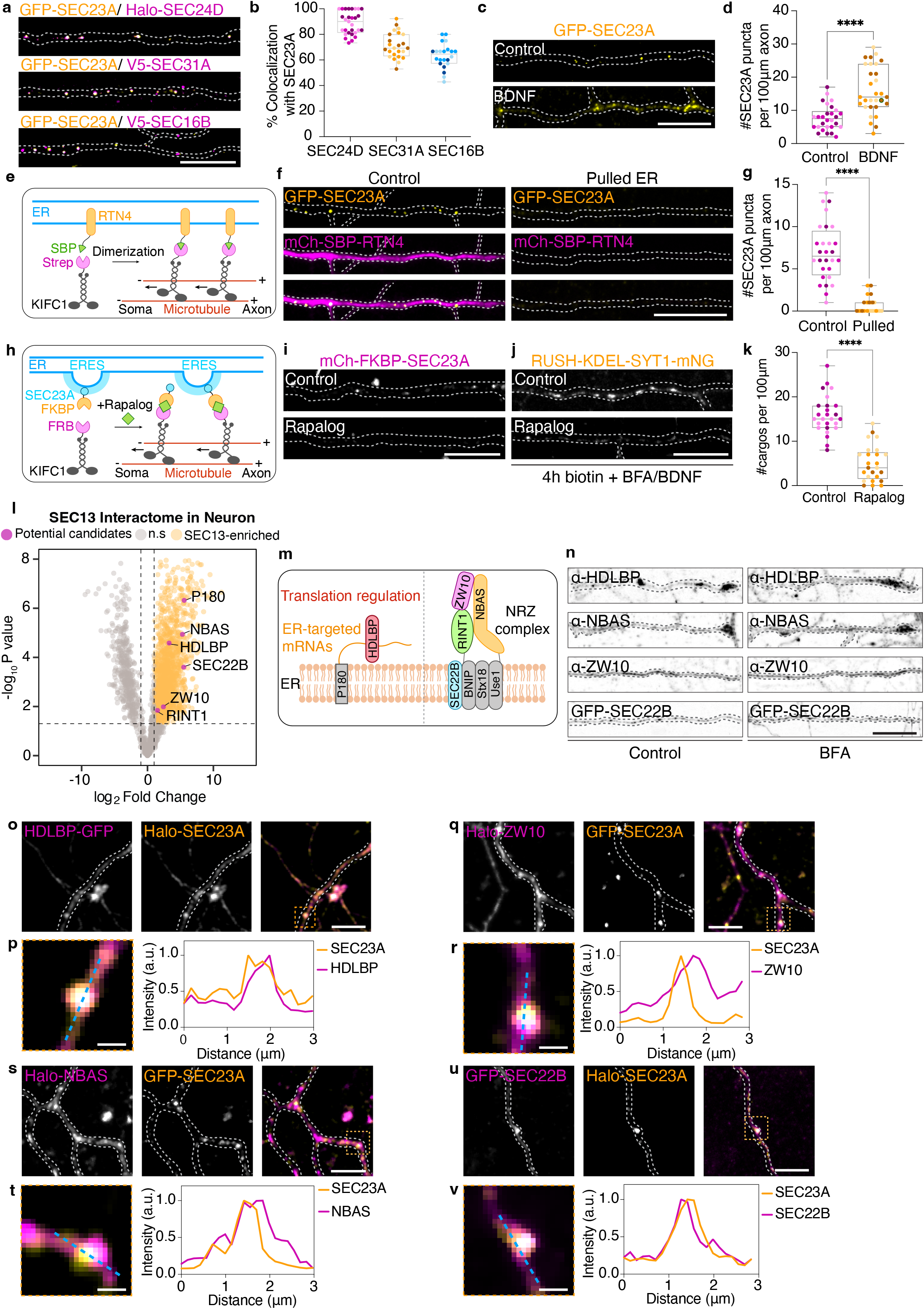
Axonal ERES are required for local secretion and ERES proxeome identifies key candidates linking local translation and secretion. **a**, Representative images of DIV9-10 axons expressing SEC23A together with SEC24D, SEC31A or SEC16B. **b**, Quantification of colocalization between SEC23A and SEC24D, SEC31A or SEC16B (N=4, SEC24D, N=3, SEC31A and SEC16B). **c**, Distribution of SEC23A in the axon in control BSA and after 30 min BDNF stimulation. **d**, Quantification of SEC23A puncta number per 100 μm axon (N=3). **e**, Schematic showing the Strep-SBP heterodimerization system using Strep-KIFC1 and SBP-RTN4 to relocate axonal ER to the soma. **f**, Representative images of SEC23A and RTN4 in DIV10 neurons in the absence (control) or presence of KIFC1 motor (pulled). **g**, Quantification of SEC23A number from conditions in **e** in 100 μm axons (N=3). **h**, Schematic showing the FKBP-FRB heterodimerization system using KIFC1-FRB and FKBP-SEC23A to relocate axonal ERES to the soma. **i**, Representative images of SEC23A treated with ethanol (vehicle, control) or Rapalog for 16h. **j**, Representative images of RUSH-SYT1 cargoes in neurons treated as in **i** for 16h, before biotin addition for 4h in the presence of BFA and BDNF. **k**, Quantification of RUSH-SYT1 cargo number from conditions in **j** in 100 μm axons (N=3). **l**, Volcano plot showing the interactome of SEC13 in neurons. Proteins that are enriched near SEC13 are in yellow and potential key candidates are highlighted in magenta. **m**, Schematic showing the localization and interactions of potential candidates at the ER membrane. **n**, Distribution of endogenous HDLBP, NBAS and ZW10 and low expression of SEC22B in axons from neurons treated or not with BFA. **o-v**, Localization and enrichment of HDLBP (**o**), ZW10 (**q**), NBAS (**s**) and SEC22B (**u**) near SEC23A in DIV10 axons. Magnifications of merged images and intensity profile lines from magnified merged images are shown in **p, r, t** and **v**. Data are presented as box-and-whisker plots in **b, d, g**, and **k**. Individual data points each represent a neuron, and each color per bar represents an independent experiment. ****p<0.0001 comparing conditions using unpaired t-test in **d, g** and **k**. Scale bars represent 10 μm in **a, c, f, i, j**, and **n**, 5 μm in **o, q, s**, and **u** and 1 μm in magnified images in **p, r, t** and **v**. See also Supplementary Figs **6-8** and Supplementary Video **4**.

We then assessed the requirement of axonal ERES for local ER exit of TMPs. For this, we employed two heterodimerization systems, the rapalog-induced FKBP/FRB and the Strep/SBP systems^43,37^ to relocate ERES components away from the axon by coupling them to the KIFC1 motor. Both tools efficiently removed axonal ERES components from the axon, without affecting axonal ER abundance (**Fig. 3h, i; Supplementary Fig. 7a-g**). We found that removal of these components from the axon before biotin addition significantly reduced the number of RUSH-SYT1 Golgi-bypassing cargoes, indicating that axonal ERES mediate TMP exit from the ER (**Fig. 3j, k**). Similar results were observed when using a SAR1B-H79G mutant, which directly inhibits ERES complex function^44,45^ (**Supplementary Fig. 7h-j**).

Since we find an important role of axonal ERES in local TMP secretion, we wondered if we could identify candidate proteins linking ERES to TMP synthesis and/or to PM delivery. We used APEX2-based proximity labeling proteomics^46^ to study the interactome of neuronal ERES using SEC13 as a bait (**Supplementary Fig. 8a**). SEC13-dependent proximal biotinylation was validated by imaging and Western blot (**Supplementary Fig. 8b, c**). In our mass spec data, we found known interactors of SEC13, including other COPII components and ER and Golgi proteins (**Supplementary Fig. 8d, e**). Moreover, we found several interesting proteins enriched near SEC13, which could have potential roles at different stages of the unconventional secretion pathway (**Fig. 3l, m**). Among these candidates, HDLBP stood out as particularly interesting for various reasons. HDLBP is an RNA-binding protein recently found to predominantly bind to ER-targeted mRNAs, promoting synthesis of transmembrane and secretory proteins^47^. Intriguingly, HDLBP interacts with the ER-associated P180/RRBP1 protein^48^, which we previously found to be enriched along the axon and involved in axonal translation (**Fig. 3m**)^17,37^. This suggests a possible link between axonal translation and secretion, in contrast to the conventional secretion in the soma, where these processes are spatially segregated: translation occurs at the rough ER and secretion occurs at the transitional ER via ERES^49,50^. Another interesting set of candidates enriched in our ERES proteome was the NRZ tethering complex (NBAS, RINT1 and ZW10) and the SNARE SEC22B, which could mediate TMP exit from the axonal ER and/or fusion with the PM. The NRZ complex is tethered to the ER via its interaction with SEC22B^51^ (**Fig. 3m**). Interestingly, recent work has also linked both the NRZ complex and SEC22B to ERES-dependent secretion in non-neuronal cells^52-54^.

We first examined whether these proteins localize to the axon. We found them present in the axonal shaft, in axonal branching points and axonal tips (**Fig. 3n; Supplementary Fig. 8f**). The distribution of all candidate proteins remained unchanged after BFA treatment (**Fig. 3n**). To determine whether these candidate proteins are in proximity to ERES, we visualized HDLBP, ZW10, NBAS and SEC22B together with SEC23A. Although these proteins broadly localize to the axonal ER (HDLBP and SEC22B) or have diffusive distribution along the axonal cytoplasm (ZW10 and NBAS), we also observed their local enrichment near axonal ERES (**Fig. 3o-v**).

These results show that axonal ERES components are associated to the axonal ER and required for unconventional TMP secretion. Moreover, we identified candidate proteins possibly linking axonal ERES to TMP translation and secretion to the PM.

### Feedback mechanism regulating local translation and secretion at axonal ERES

Since the ER-translation regulator HLDBP was found near axonal ERES, we wondered whether TMP mRNAs can also be found near ERES. To assess this, we co-expressed the ERES protein SEC23A and a SYT1 mRNA reporter containing PP7 RNA stem loops. Live-cell imaging revealed similar dynamics of ERES and SYT1 mRNAs in the axon, with both showing predominantly stable movements (**Fig. 4a**). Interestingly, a subset of SYT1 mRNAs localized near axonal ERES, with ± 60% of SEC23A puncta lying in proximity to SYT1 mRNA and ±40% of SYT1 mRNA particles positioned near SEC23A (**Fig. 4b**). Next, we visualized SYT1 translation sites with the Puro-PLA system and their co-distribution with SEC23A in the axon. We observed instances of close proximity between translation sites and SEC23A in axon shafts, as well as at branching points (**Fig. 4c**). To further confirm this, we employed the cytERM-SunTag system to visualize ER-targeted translation^55^, where the ER-anchoring sequence cytERM is fused to 24 tandem GCN4 repeats in its N-terminal. Co-expressed scFv-GFP antibodies bind these epitopes as they emerge from the ribosome during translation, amplifying the signal up to 24-fold for bright detection of active translation (**Fig. 4d**). Using this method, we also detected translation sites near axonal ERES (**Fig. 4e**).

**Fig. 4:**
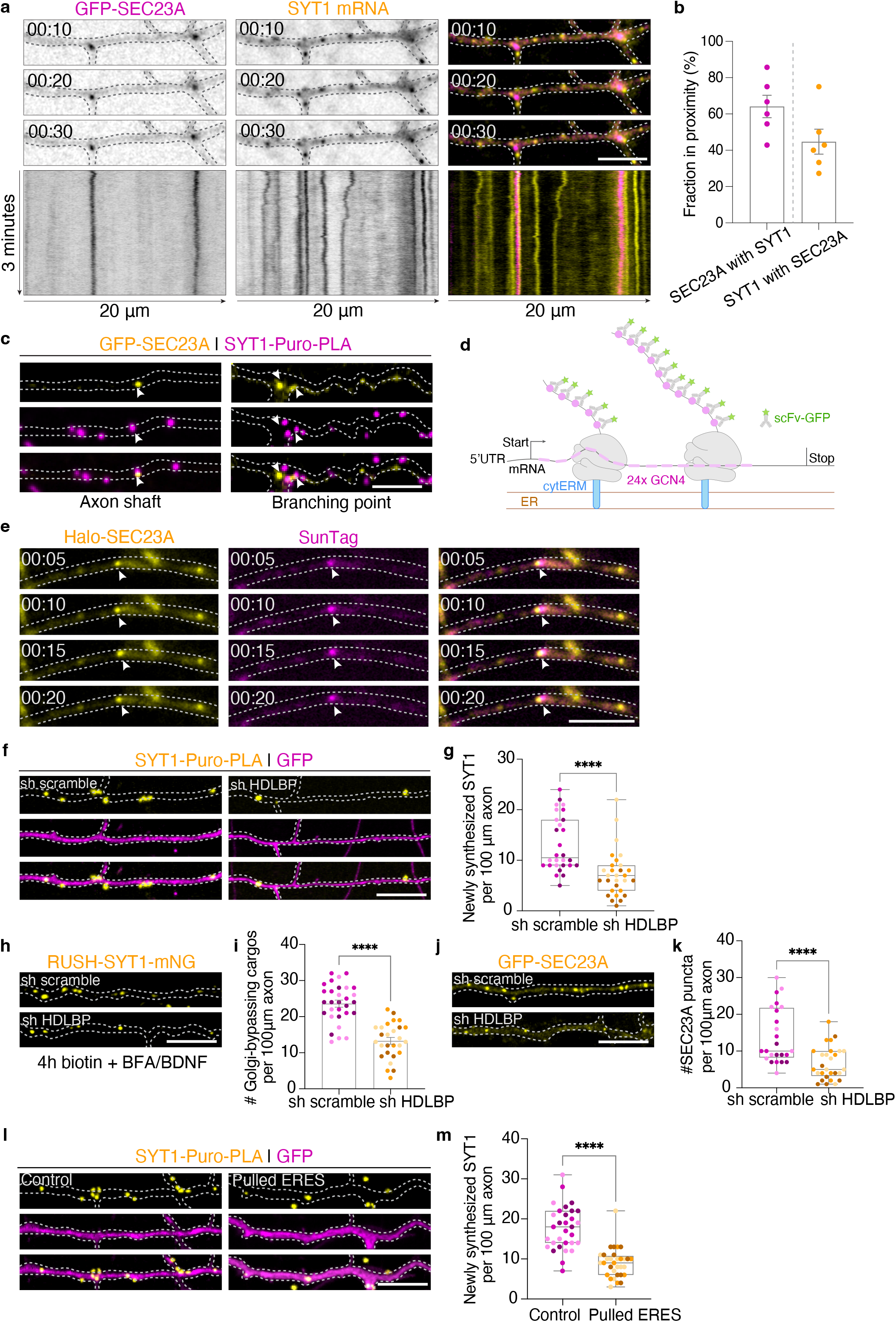
Feedback mechanism regulating translation and secretion at axonal ERES. **a**, Time series showing localization and co-movements of SEC23A and SYT1 mRNA in the axon. Images were acquired at 1-second intervals for 3 minutes. **b**, Quantification of the fraction of SEC23A in proximity to SYT1 mRNA and of SYT1 mRNA in proximity to SEC23A (N=1). **c**, Representative images of SYT1 translation sites with Puro-PLA assay in proximity to SEC23A in the axon. **d**, Schematic representation of the cytERM-SunTag system to visualize ER-targeted translation. **e**, Time series showing localization and co-movements of SEC23A and SunTag translation signals in the axon. Images were acquired at 1-second intervals for 3 minutes. **f**, Representative images of SYT1 translation sites using the Puro-PLA assay in axons of DIV9-10 neurons expressing a scramble shRNA (control) or shRNA targeting HDLBP. **g**, Quantification of newly synthesized SYT1 number per 100 μm axon (N=3). **h, i**, Representative images of DIV11 neurons expressing RUSH-SYT1, together with shRNA scramble or shRNA targeting HDLBP. Neurons were treated with BFA, biotin and BDNF for 4h (**h**). Quantification of Golgi-bypassing cargo number for conditions in **h** in 100 μm axon (**i**) (N=3). **j, k**, Representative images of DIV10-11 neurons expressing GFP-SEC23A together with shRNA scramble or shRNA targeting HDLBP (**j**). Quantification of SEC23A puncta in 100 μm axon (**k**) (N=3). **l, m**, Representative images of SYT1 translation sites using the Puro-PLA assay in axons of DIV10 neurons in control condition or after ERES pulling (**l**). Quantification of newly synthesized SYT1puncta in 100 μm axon (**m**). Data are presented as box-and-whisker plots in **g, k** and **m** or as mean values ± SEM in **b** and **i**. Individual data points each represent a neuron, and each color per condition represents an independent experiment. ****p<0.0001 comparing conditions with control using Mann-Whitney test in **g** and **m** and unpaired t-test in **i** and **k**. Scale bars represent 5 μm in **a, c** and **e** and 10 μm in **f, h, j** and **l**. See also Supplementary Fig. **9**.

Since HDLBP has been shown to regulate translation of TMPs in non-neuronal cells^47^, we wondered if HDLBP is also involved in axonal TMP synthesis. We performed knockdown of HDLBP in combination with SYT1 Puro-PLA assay to visualize axonal SYT1 translation (**Fig. 4f, g; Supplementary Fig. 9a**). Compared to our sh scramble control, HDLBP knockdown led to reduced SYT1 translation, from ±13 translation sites/100 μm in scramble shRNA control to ±7 sites/100 μm in HDLBP knockdown condition (**Fig. 4f, g**). We next examined if HDLBP-dependent translation has an impact on cargo exit from the ER. To investigate this, we visualized RUSH-SYT1-mNG unconventional secretion in the presence of scramble or HDLBP shRNA. HDLBP knockdown impaired unconventional cargo secretion in the axon (**Fig. 4h, i**). Notably, in our experimental setup, we simultaneously co-expressed the RUSH-SYT1 cargoes and HDLBP shRNA, resulting in RUSH cargoes being produced at the ER prior to HDLBP knockdown (**Supplementary Fig. 9e, f**). The reduction in the number of Golgi-bypassing cargoes exiting the axonal ER in knockdown condition therefore suggests a role of HDLBP in coupling TMP synthesis to ER exit beyond translation regulation. To assess the involvement of HDLBP in coupling TMP synthesis with secretion, we looked at ERES and found that HDLBP knockdown caused a reduction in the number of ERES puncta in the axon, from ±14 puncta to ±6 puncta (**Fig. 4j, k**). Reciprocally, we asked whether disrupting axonal ERES affects local TMP synthesis. To study this, we removed axonal ERES components and looked at SYT1 translation, which decreased from ±18 to ±9 puncta (**Fig. 4l, m**). Together, these bidirectional effects reveal a feedback loop between axonal TMP synthesis and secretion at axonal ERES, possibly coordinated by HDLBP.

### The NRZ-SEC22B tethering complex promotes local TMP unconventional secretion, dependent on ER-PM contacts

We also identified the NRZ-SEC22B tethering complex in our neuronal ERES proteome, which could bridge ER exit and/or PM insertion of axonally synthesized TMPs. The SNARE protein SEC22B cycles between the ER and acceptor membranes to drive membrane fusion in its unassembled/non-tethered form^53,54^. On the other hand, the SEC22B-interacting NRZ tethering complex has been suggested to recruit acceptor membranes to ERES for cargo exit^52,54^. Intriguingly, SEC22B has an additional non-fusogenic role at ER-PM contact sites ^56,57^. To further study the role of these candidate proteins in axonal TMP ER exit and PM fusion, we first used live-cell imaging to look at the distribution and dynamics of SEC22B with the NRZ complex, and with neosynthesized SYT1 cargoes. When visualized together with either ZW10, NBAS or RUSH-SYT1, a portion of SEC22B puncta colocalized with these proteins and had similar short-range, bi-directional movements in the axon, suggesting a possible involvement of SEC22B and the NRZ tethering complex in local cargo secretion (**Fig. 5a-c; Supplementary Video 5**). Given the role of SEC22B in bridging ER-PM contact sites for axonal membrane expansion^56^, we hypothesized that Golgi-bypassing cargoes might be locally sorted to the PM, proximal to these sites. Indeed, using the GFP-MAPPER tool to visualize ER-PM contact sites^58-60^, we observed cargoes located near these sites (**Fig. 5d**).

**Fig. 5:**
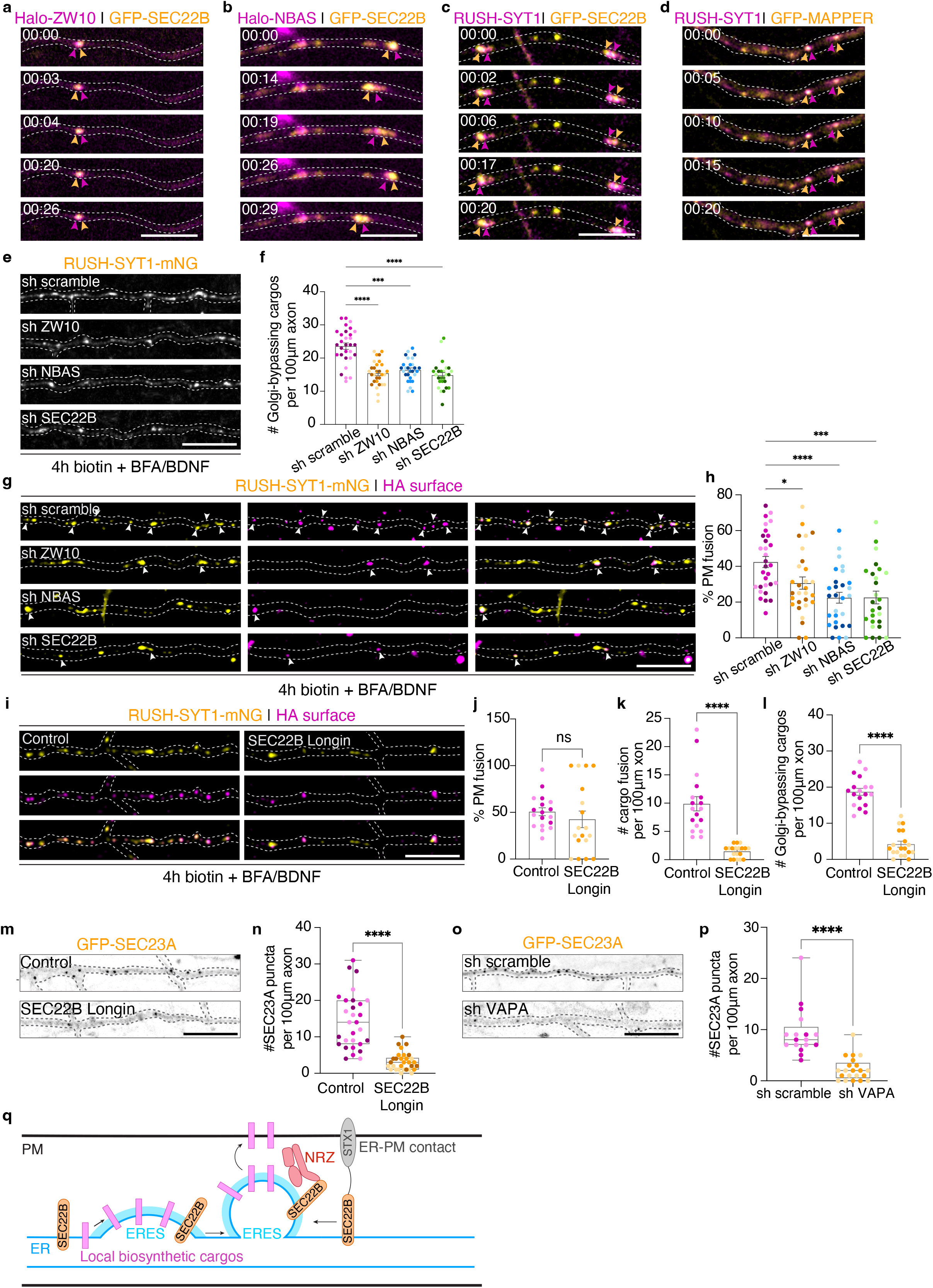
SEC22B and the NRZ tethering complex promotes local TMP secretion, dependent on ER-PM contact sites. **a-d**, Time series showing colocalization and co-movements of SEC22B with ZW10 (**a**), NBAS (**b**) and Golgi-bypassing RUSH-SYT cargoes after BFA treatment for 30 min (**c**). Time series showing colocalization and co-movements of RUSH-SYT1 cargoes after BFA treatment for 30 min with the ER-PM marker GFP-MAPPER (**d**). Images were acquired at 1-second intervals for 3 minutes. Arrowheads point to the colocalized regions. **e**, Representative images of DIV11 neurons expressing RUSH-SYT1, together with shRNA scramble or shRNAs targeting ZW10, NBAS or SEC22B. Neurons were treated with BFA, biotin and BDNF for 4h. **f**, Quantification of Golgi-bypassing cargo number for conditions in **e** in 100 μm axon (N=3). **g**, Representative images of DIV11 neurons transfected as in (**e**), and live cell surface labeled for HA-tag incorporated to the luminal/extracellular side of RUSH-SYT1. Arrowheads point to RUSH-SYT1 cargoes that are also positive for HA labeling. **h**, Quantification of the percentage of Golgi-bypassing cargoes fusing with the PM (N=3). **i-l**, Representative images of DIV9 neurons expressing RUSH-SYT1 alone or together with SEC22B Longin domain. Neurons were treated with BFA, biotin and BDNF for 4h (**i**). Quantification of the percentage of Golgi-bypassing cargoes fusing with the PM for conditions in **i** (**j**). Quantification of total number of cargoes fusing with the PM in 100 μm axon (**k**). Quantification of total number of Golgi-bypassing cargo in 100 μm axon (**l**) (N=2). **m, n**, Representative images of DIV9 neurons expressing SEC23A alone, or together with SEC22B Longin domain (**m**). Quantification of number of SEC23A puncta in 100 μm axon (**n**) (N=3). **o, p**, Representative images of DIV10 neurons expressing SEC23A together with sh scramble or shRNA targeting VAPA (**o**). Quantification of number of SEC23A puncta in the axon in 100 μm axon (**p**) (N=2). **q**, Model schematic for the role of NRZ-SEC22B tethering complex in cargo exit and delivery to the PM. Data are presented as mean values ± SEM in **f, h, j, k** and **l** or as box-and-whisker plots in **n** and **p**. Individual data points each represent a neuron, and each color per condition represents an independent experiment. ns= non-significant, p=0.4174 comparing conditions using unpaired t-test in **j**, *p<0.05, ***p<0.001, ****p<0.0001 comparing conditions with control using Kruskal-Wallis test followed by Dunn’s multiple comparisons test in **f**, ordinary one-way ANOVA test followed by Dunnett’s multiple comparisons test in **h**, Mann-Whitney test in **k, n** and **p**, or unpaired t-test in **l**. Scale bars represent 5 μm in **a, b, c** and **d**, and 10 μm in **e, g, i, m** and **o**. See also Supplementary Fig. **9** and Supplementary Video **5**.

Since SEC22B and the NRZ complex co-distribute with each other (**Fig. 5a, b**), with ERES (**Fig. 3q-v**) and Golgi bypassing cargoes (**Fig. 5c**) along the axon, we wondered if they could play a role in unconventional secretion of TMPs along the axon. Thus, we transfected neurons with shRNAs targeting SEC22B, ZW10, and NBAS together with RUSH-SYT1-mNG (**Fig. 5e, f; Supplementary Fig. 9b-d**). We observed a similar reduction in the number of SYT1 Golgi-bypassing cargoes following knockdown of SEC22B and NRZ complex proteins, compared with control (**Fig. 5e, f**). Intriguingly, we noted that knockdown of SEC22B and the NRZ complex does not seem to affect somatic conventional secretion. We observed similar dynamics in cargo sorting from ER to Golgi after 30min ER release (**Supplementary Fig. 9e, f**). This suggests that these proteins have a more critical function in unconventional secretion in the axon. We also assessed the ability of RUSH-SYT1 cargoes to fuse with the PM upon the different knockdown conditions. NRZ complex and SEC22B knockdown resulted in a reduced percentage of unconventional cargoes reaching the PM (30% of cargoes for ZW10, 22.4% for NBAS, and 22.5% for SEC22B, with respect to 42% in control) (**Fig. 5g, h**).

To gain more mechanistic insights into the role of SEC22B’s tethering function at ER-PM contact sites in the sorting of neosynthesized TMP to the PM, we employed a dominant negative approach. The SEC22B Longin domain is essential for recruiting E-SYT1 to stabilize the non-fusogenic SEC22B-STX1 complex at ER-PM contact sites. Expression of the Longin domain, which competes with endogenous SEC22B at ER-PM contact sites, leads to impaired PM expansion^56,57^. We observed that expression of the SEC22B Longin domain (1-110aa), unexpectedly, caused impaired TMP secretion due to a drastic reduction in ERES number along the axon (**Fig. 5i-n**). This result led us to wonder whether ER-PM contact sites broadly modulate local unconventional TMP secretion by regulating axonal ERES number. Similar to overexpression of SEC22B Longin domain, knockdown of the ER tethering protein VAPA, involved in ER-PM contact site assembly^61^, reduced ERES number along the axon (**Fig. 5o, p**).

Together, these results show that the SEC22B-NRZ complex is required for local TMP sorting from the ER to the PM, and that disruption of ER-PM contacts affects ERES assembly. These results, together with previous findings about SEC22B-NRZ interactions, leads us to propose a model where the NRZ complex is recruited to ERES by SEC22B to stabilize the ER SNARE and recruit acceptor membranes for cargo exit at ERES. Within the narrow diameter of the axon, this acceptor could be the PM. Via this mechanism, this SEC22B-NRZ complex could then locally sort cargoes to the PM proximal to SEC22B-STX1 ER-PM contact sites (**Fig. 5q**).

### Unconventional TMP secretion supports axon growth and synaptic bouton maturation during axonal development

TMPs such as the adhesion molecules and synaptic proteins shown here to undergo axonal translation and unconventional secretion, play essential roles in axon development and maintenance^18,62-64^. Therefore, we wondered if perturbation of local unconventional secretion results in defects in axon development and maintenance. To study this, we first used the Strep/SBP heterodimerization tool for long-term local removal of axonal ERES components (**Supplementary Fig. 7a-g**). We first investigated whether unconventional axonal secretion is required during early stages of development. Previous studies have depleted the ERES component SAR1 but found conflicting results regarding its contribution to axon growth^40,41^. Here, we transfected neurons at DIV2 with our axonal ERES removal tool and fixed them at DIV4. We observed that by removing ERES components from the axon, the primary axon length was significantly reduced, from ± 400 μm to ± 305 μm (**Fig. 6a, b**). We also found that the total axon arborization was reduced, from ± 1167 μm to ± 634 μm, indicating a decrease in overall axon growth (**Fig. 6a, c**). These results support a local role of ERES in axon development.

**Fig. 6:**
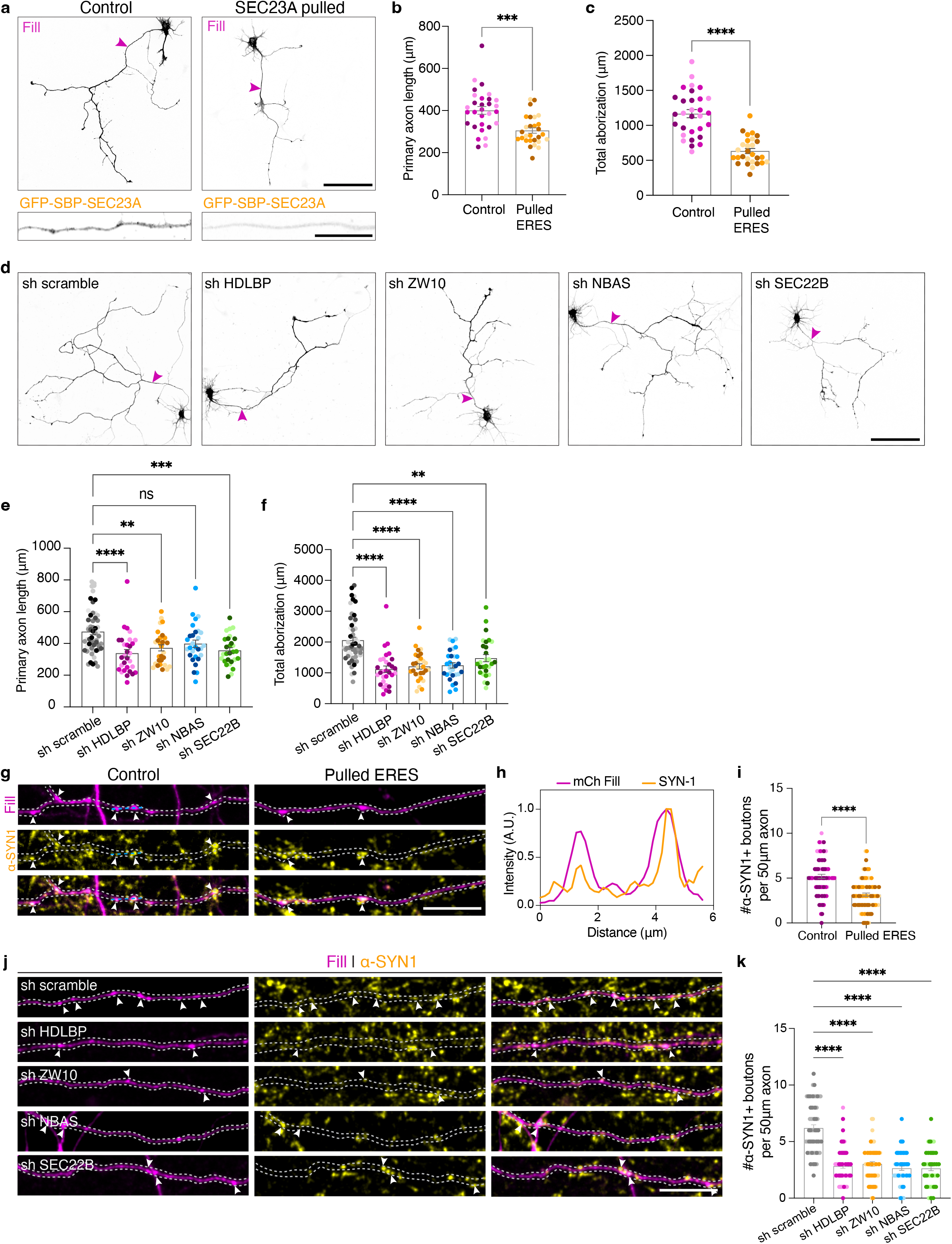
Unconventional secretion of TMPs promotes axon growth and pre-synaptic bouton assembly. **a-c**, Representative images of DIV4 neurons transfected with a fill and SBP-SEC23A (control) or with Strep-KIFC1 (SEC23A pulled) (**a**). Quantification of primary axon length (**b**) and total axon arborization (**c**). (N=3) Arrowheads point to axons. **d-f**, Representative images of DIV4 neurons transfected with a fill and a scramble shRNA or shRNAs targeting HDLBP, ZW10, NBAS or SEC22B (**d**). Quantification of primary axon length (**e**) and total axon arborization (**f**). (N=3). Arrowheads point to axons. **g-i**, Representative images of DIV17 neurons transfected with a fill and SBP-SEC23A (control) or with Strep-KIFC1 (SEC23A pulled). Staining for endogenous Synapsin-I (SYN1) was performed to label mature pre-synaptic boutons. Arrowheads point to boutons positive for SYN1 (**g**). Intensity profile lines from highlighted boutons in **g** (**h**). Quantification of mature bouton number in 50 μm axon length (**i**) (N=3). **j, k**, Representative images of DIV17 neurons transfected with a fill and a scramble shRNA or shRNAs targeting HDLBP, ZW10, NBAS, or SEC22B and stained for endogenous SYN1 (**j**). Quantification of mature bouton number in 50 μm axon length (**k**) (N=2). Data are presented as mean values ± SEM in **b, c, e, f, i** and **k**. Individual data points each represent a neuron in **b, c, e** and **f**, or an axon segment in **i** and **k**, and each color per condition represents an independent experiment. ns = non-significant (p=0.0934), **p < 0.01, ***p < 0.001, ****p<0.0001 comparing conditions to control using Mann-Whitney test in **b**, unpaired t-test in **c** and **i**, Kruskal-Wallis test followed by Dunn’s multiple comparisons test in **e, f, k**. Scale bars represent 100 μm in overview images in **a, d**, 20 μm in axon crops in **a** and 10 μm in **g** and **j**. See also Supplementary Fig. **7**.

To study the role of identified proteins involved in unconventional secretion on axon development, we transfected DIV1 neurons with a scramble shRNA control, or shRNAs targeting HDLBP, ZW10, NBAS, or SEC22B. Similar to what was observed for axonal ERES removal, knockdowns of these proteins resulted in shorter primary axons (**Fig. 6d, e**) and reduced total axon arborization (**Fig. 6d, f**). This suggests that unconventional secretion of neosynthesized TMPs from axonal ERES, regulated by HDLBP and NRZ-SEC22B complex, is essential for axon growth.

During neurodevelopment, after axon elongation and branching, the assembly of presynaptic boutons and eventually functional synapses is of great importance. Synapse formation and maintenance is a complex process that requires many TMPs to be recruited and assembled in pre- and post-synapses^65-67^. Local protein synthesis is critical for the development and function of these synapses^68^. Therefore, we wondered whether unconventional secretion of newly synthesized proteins in the axon plays a role in pre-synaptic bouton assembly. We first performed local removal of axonal ERES components using our Strep-SBP heterodimerization tool (**Supplementary Fig. 7a-g**). Neurons were transfected on DIV14 with SEC23A-SBP and KIFC1-Strep heterodimerization system. Co-expression of a fluorescent protein (fill) was used to identify pre-synaptic boutons in transfected axons. DIV17-18 neurons were then fixed and stained for the peripherally synaptic vesicle-associated protein Synapsin-I (SYN1) to identify mature presynaptic boutons (**Fig. 6g, h**). Removal of ERES components led to a lower number of boutons that were positive for SYN1, from ±5.2 boutons to ±3.1 boutons per 50 μm axon, suggesting that local secretion is required for synapse assembly (**Fig. 6g, i**).

Next, we studied the role of ERES-associated proteins involved in unconventional secretion in pre-synaptic bouton assembly. We transfected DIV12-14 neurons with a fluorescent fill and a scramble shRNA control, or shRNAs targeting HDLBP, ZW10, NBAS, or SEC22B. Neurons were then fixed at DIV16-18 and stained for SYN1, as described above. Knockdown of these proteins led to similar results as for the removal of axonal ERES components (**Fig. 6j, k**).

Together, these results indicate that the local unconventional secretion route, described in this study, plays a role in axon development and pre-synaptic bouton assembly.

## Discussion

For many years, it has been believed that the supply of TMPs to the axon strictly depends on their production and trafficking from the soma. In this study, we show that TMPs can be locally translated and secreted to the axonal PM. Exit of TMPs from the axonal ER requires ERES but not Golgi-derived organelles. We identified the ER-translation regulator HDLBP, and the membrane tethering NRZ-SEC22B complex associated with axonal ERES, which are involved in different steps of axonal cargo synthesis, ER exit and delivery to the PM (**Fig.7**). HDLBP coordinates local translation and secretion at local ERES via a bi-directional feedback mechanism, while SEC22B and the NRZ complex promote cargo exit from the ER and delivery to the axonal PM, dependent on ER-PM interactions. This axonal unconventional protein secretion, which is increased upon neuronal stimulation, plays a key role during axon growth and pre-synaptic bouton assembly (**Fig. 7**).

**Fig. 7:**
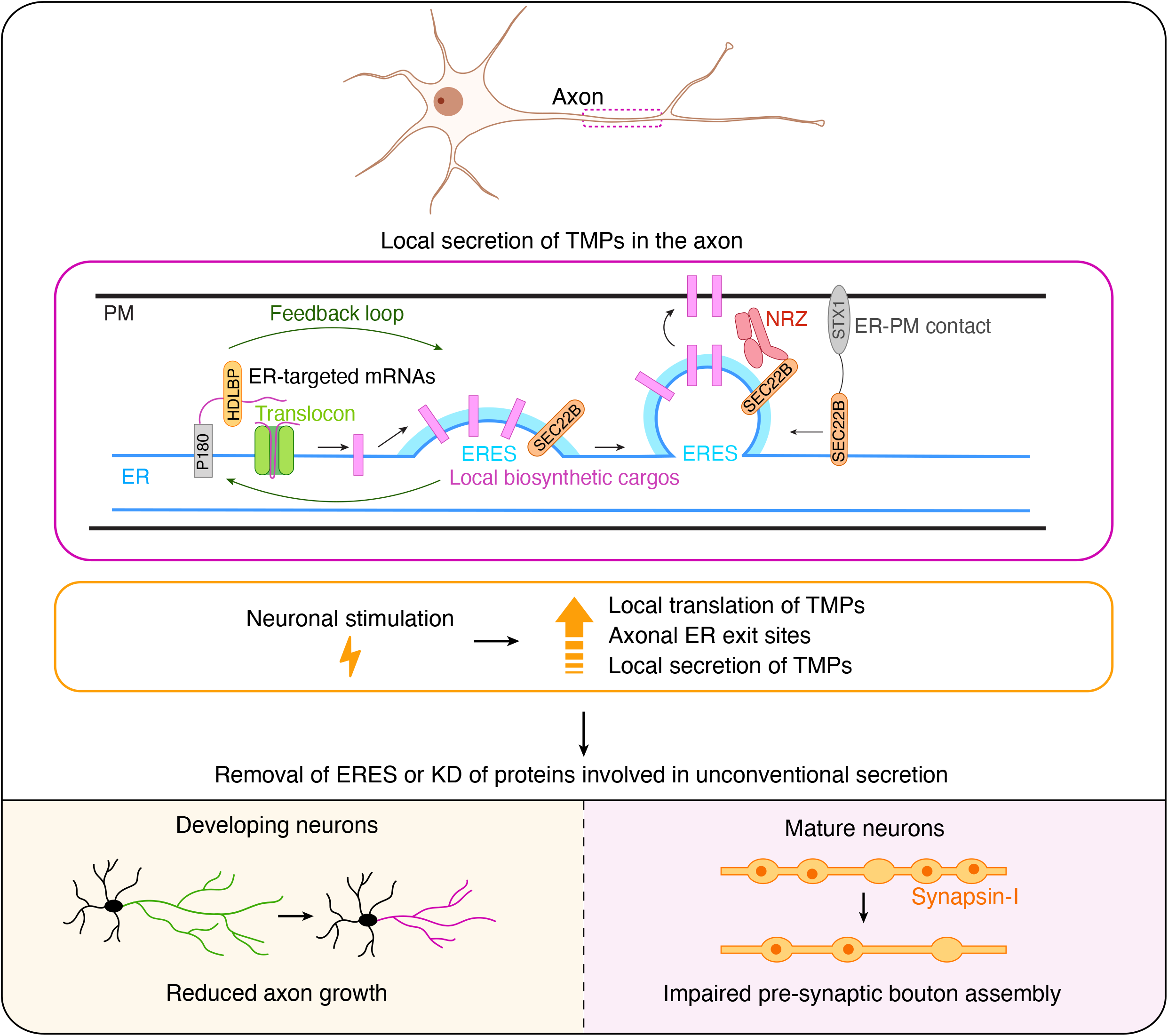
Coupling of local translation and secretion in the axon. Locally translated TMPs exit the ER via axonal ERES, followed by their delivery to the PM. This process is dependent on the translation regulator HDLBP and NRZ-SEC22B tethering complex, which are involved in different steps of axonal cargo synthesis, ER exit and delivery to the PM. HDLBP coordinates local translation and secretion at local ERES via a bi-directional feedback mechanism, while SEC22B and the NRZ complex promote cargo exit from the ER and delivery to the axonal PM, regulated by ER-PM interactions. This axonal unconventional protein secretion, which is induced upon neuronal stimulation, plays a key role during axon growth and pre-synaptic bouton assembly.

Consistent with several transcriptomic and translatomic studies in isolated axons^69,8,11^, we found that TMP mRNAs are present and translated in axons, and their translation increases upon neuronal stimulation. How do these locally translated TMPs exit the ER and reach the axonal PM? Notably, we did not observe Golgi-related compartments along the axon, in contrast to their contribution to local conventional ER to PM secretion in dendrites^32,30,31,70^. This led us to wonder whether axonally translated TMPs have the capability to exit the ER upon ER-Golgi trafficking dysfunction. We found that a subset of TMPs was still able to exit the ER and reach the axonal PM upon BFA treatment. Importantly, axonal TMPs were locally and unconventionally secreted from the axonal ER as shown in our experiments in desomatized axons. Although ERES components have been previously found along the axon^40,41^, it remained unknown whether axonal ERES components contribute to local unconventional TMP secretion. We found that these ERES components form a complex stably associated with the axonal ER. Importantly, axonal removal of ERES prevented TMP exit from the axonal ER, and neuronal stimulation increased ERES number to accommodate increased neosynthesized TMP secretion. The mechanism behind ERES number increase upon neuronal stimulation remains unknown. It is possible that cytosolic ERES components are already distributed along the axon, or locally synthesized, to assemble on the ER membrane for fast and local delivery of newly synthesized TMPs to the PM. The fact that these Golgi-bypassing cargoes can reach the axonal PM raises the question whether TMPs, in a core-glycosylated state, are functional. Interestingly, previous studies have shown that many core-glycosylated dendritic and axonal TMPs are sorted to the PM, without being processed at the GA or Golgi-derived compartments^71-73^. These TMPs are delivered to the PM to allow for fast response to local needs. They maintain their functionality at the PM, although with potentially higher turnover rate or a change in their activity at the PM surface^73,74^.

Given that axonal length and area far exceed those of the soma, it is unlikely that the molecular machineries for translation, ER exit, and secretion are spatially segregated. Our ERES proximity labeling proteomic analysis in primary rat neurons revealed a set of candidate proteins localized near ERES, which could coordinate translation and secretory processes within the axon. One such protein is HDLBP, an RNA-binding protein that selectively binds to ER-targeted mRNAs^47^ and interacts with the ER-resident and axon-enriched protein P180^37,48,17^. We previously found that P180 binds TMP mRNAs and is essential for axonal translation regulation^17^. Here, we found that HDLBP is present along the axon and is in close proximity to ERES. Consistently, we also observed mRNAs and local translation of TMPs near ERES. Of note, the role of HDLBP has been only studied in cell lines. Here, we showed that HDLBP knockdown significantly reduced SYT1 synthesis in the axon, further supporting the existence of an axonal ER-associated translation machinery responsible for local TMP production^17^. Unexpectedly, HDLBP knockdown also impaired pre-existing cargo exit from the axonal, but not somatic, ER. This effect was associated with impaired ERES assembly, suggesting coupling of TMP synthesis and ER exit along the axon. Conversely, removal of axonal ERES reduced SYT1 translation sites in the axon. This points towards a feedback regulation between axonal TMP synthesis and secretion at axonal ERES, possibly coordinated by HDLBP.

Having established this coupling between local translation and ER export, we next asked how these cargoes subsequently reach the PM. Vesicle tethering proteins are critical mediators of membrane fusion events between donor and acceptor compartments^75,76^. In our proteomic analysis, we identified components of the NRZ tethering complex as well as the SNARE SEC22B. We observed these components to be distributed along the axon and found they were required for cargo exit and targeting to the axonal PM. The NRZ complex, composed of NBAS, RINT1 and ZW10, is classically linked to Golgi-to-ER retrograde transport, where it stabilizes the ER-resident SNARE complex containing SEC22B, BNIP, STX18, and USE1 to mediate fusion of COPI-coated vesicles at the ER^51^. Moreover, SEC22B is also implicated in broader vesicular trafficking^54^ and in ER-PM contact formation and maintenance^56,57^. However, we did not observe Golgi-derived compartments nor COPI components along the axon, suggesting an unconventional role for the NRZ-SEC22B complex in cargo sorting to the axonal PM. Indeed, recent work has linked NRZ to anterograde canonical^52^ and non-canonical secretory pathways^77^. The NRZ complex has been shown to tether and recruit ERGIC membrane to ERES^52^, creating a tunnel between the ER and Golgi for collagen export^78^. Consistent with this, recent evidence suggests that ERES/COPII components remain at the neck of cargo-decorated ER tubular network reaching the acceptor membranes^79^. Building on these findings, in the narrow confines of the axon where Golgi-related organelles are absent, SEC22B, together with the NRZ complex could facilitate a similar local mechanism to sort biosynthetic cargo from the ER to the PM. In this model, SEC22B and the NRZ complex can tether and stabilize the ER to the PM, upon which cargoes would be delivered along sorting intermediates extending from ERES towards the PM, while COPII components remain at the ERES neck. Alternatively, ER-derived vesicles budding from ERES could deliver cargoes to the PM. In either case, this ER to PM sorting must be local, as we did not observe cargoes nor the NRZ-SEC22B complex undergoing long-range transport along the axon. Interestingly, we observed neosynthesized cargoes next to ER-PM contact sites. Surprisingly, we found that expression of SEC22B Longin domain, which is involved in the non-fusogenic role of SEC22B in ER-PM assembly^56^, impaired cargo secretion by disrupting ERES assembly. Similarly, disrupting ER-PM tether VAPA resulted in comparable defects. Together, these results indicate that ER-PM contact sites regulate TMP transfer from the ER to PM. Although ER-PM contact sites are typically associated with ion and lipid exchange, such exchanges could also be essential for regulated local unconventional secretion. Indeed, calcium and PI4P exchange occurs at ER-PM contacts^80,81^ and are important for ERES function^82,83^.

We further studied the role of axonal ERES and its coupling to translation and secretion machineries in axon growth and pre-synaptic bouton assembly. During early development, axons navigate through their environment and respond to extracellular cues through TMPs including cell adhesion molecules, receptors and ion channels at axonal growth cones or tips^5,62,63,84^. These proteins are also essential later for synapse formation and maintenance, where pre-and post-synaptic assembly is tightly regulated^18,64,85^. Here, we show that removal of axonal ERES components or knockdown of HDLBP or NRZ-SEC22B complex at different developmental stages reduced primary axon length, total arborization, and impaired pre-synaptic bouton assembly. Together, our findings support a key role for the local coupling of TMP synthesis and secretion during axonal development. They also raise new open questions for future research. For instance, how does this local axonal machinery for TMP production and secretion contribute to shaping the axonal proteome? Do local translation and secretion diversify TMP and neuron functioning? Finally, considering mutations in ER-resident proteins that alter axonal ER organization also cause Hereditary Spastic Paraplegia (HSP) and Amyotrophic Lateral Sclerosis (ALS), what is the contribution of axonal ER-associated translation and secretion defects to disease?

Altogether, our findings reveal an axon-intrinsic unconventional secretory pathway that couples local translation, ER export, and PM insertion of TMPs in CNS axons. This pathway relies on the coordinated action of local ERES with HDLBP-mediated translation and NRZ-SEC22B-regulated secretion. Its response to external cues further supports the adaptability of this pathway to meet spatiotemporal axonal needs. Its essential role across neuronal development, from axon growth to presynaptic bouton assembly, underscores the physiological importance of localized TMP synthesis and secretion for proper neuronal development and maintenance.

## Methods

### Animals

All experiments were approved by the DEC Dutch Animal Experiments Committee (Dier Experimenten Commissie), performed in line with institutional guidelines of Utrecht University, and conducted in agreement with Dutch law (Wet op de Dierproeven, 1996) and European regulations (Directive 2010/63/EU). The animal protocol has been evaluated and approved by the national CCD authority (license AVD10800202216383). Female pregnant Wistar rats were obtained from Janvier, and embryos (both genders) at E18 stage of development were used for primary cultures of hippocampal neurons. Brains from these female rats were used to obtain protein extracts. At Janvier, rats were housed in a temperature and humidity-controlled facility at a 12h/12h light cycle in cages with spruce bedding with enrichment. Pregnancy was monitored by visual examination and weighing where a significant weight gain meant gestation. Pregnant females were then shipped to the Central Laboratory Animal Research Facility of Utrecht University where they were housed for 4 days in pairs under standard laboratory conditions and received food and water ad libitum. Pregnant females were then euthanized, and embryos (E18) were harvested for primary neuronal cultures. The animals, pregnant females and embryos have not been involved in previous procedures.

### Primary neuronal cultures and transfection

Primary hippocampal neurons were prepared from embryonic day 18 rat brains from which the hippocampi were dissected, dissociated in trypsin for 15 min at 37°C and plated at a density of 100,000/well or 50,000/well (12-well plates) on coverslips coated with poly-L-lysine (37.5 μg/mL; Sigma) and Laminin (1.25 μg/mL; Roche). For proteomics, cortices were dissected and dissociated as hippocampal neurons and plated at a density of 1,000,000/well (6-well plates). For experiments in developing neurons (until DIV11), neurons were maintained in neurobasal medium (NB; Gibco) supplemented with 2% B27 (Gibco), 0.5 mM Glutamine (Gibco), 15.6 μM Glutamate (Sigma), and 1% Penicillin/Streptomycin (Gibco) and incubated under controlled temperature and CO2 conditions (37°C, 5% CO2). For experiments in mature neurons (until DIV18), medium was refreshed at DIV10 by replacing half of the medium with BrainPhys (BP) neuronal medium supplemented with 2% Neurocult SM1 neuronal supplement and 1% Pen/Strep. For all RUSH experiments, B27 was subjected to biotin removal using Zeba™ Dye and Biotin Removal Spin Columns (Invitrogen) according to manufacturer’s protocol.

Hippocampal neurons were transfected at day in vitro (DIV)1-2 (for DIV 4 experiments), or DIV7-9 (for DIV10-11 experiments) or DIV11-14 (for DIV16-18 experiments) using Lipofectamine 2000 (Invitrogen). Shortly, DNA (0.05-1.5 μg/well) was mixed with Lipofectamine 2000 (1.5-3.3 μL) in NB (Gibco, 200 μL) and incubated for 20 min at room temperature. Before transfection, half of the original medium from each well was stored (500 uL), and 200 uL was removed. For RUSH experiments, 700 uL of the original medium in each well was removed. The transfection mix was added to the neurons in the remaining NB+ medium and incubated for 1 hour at 37°C in 5% CO2. Then, 500 uL of original medium plus 500 uL of fresh NB or BP medium with supplements were added to each well. For RUSH experiments, neurons were transferred to a new plate containing 1 mL of NB RUSH medium in each well. Neurons were kept at 37°C in 5% CO2 until fixation.

### Human iPSC culture and neuronal differentiation

Human iPSCs (KOLF2.1J)^87^ were differentiated into cortical neurons (iNeurons) by induced expression of human NGN2, following a previously described protocol^88^ with minor adaptations. Briefly, KOLF2.1J iPSCs were cultured in Stemflex medium (Thermo Fisher Scientific, Cat# A3349401) in a 6-well plate coated with Matrigel (Merck, Cat# CLS354277). iPSCs were transfected with PB-TO-hNGN2 (Addgene, #172115) and a plasmid containing EF1a-transposase using Lipofectamine Stem (Thermo Fisher Scientific, Cat# STEM00001), followed by selection with 2 μg/ml puromycin to generate a polyclonal pool of iPSCs with stably integrated tet-inducible human NGN2.

Differentiation towards iNeurons was induced by plating these iPSCs in a Matrigel-coated 6-well plate in induction medium (DMEM/F12 (Thermo Fisher Scientific, Cat# 11330032) with 1x N2 supplement (Thermo Fisher Scientific, Cat# 17502048), 1x non-essential amino acids (Thermo Fisher Scientific, Cat# 11140050), 2 mM L-glutamine (Thermo Fisher Scientific, Cat# 25030081), 2 μg/ml Doxycycline (Stemcell Technologies, Cat# 72742)). After three days, the cells were dissociated with accutase (Thermo Fisher Scientific, Cat# A1110501), and plated on glass coverslips coated with poly-L-lysine (Sigma) in maturation BP medium (Stemcell Technologies, Cat# 05790) supplemented with 1x B27 supplement (Gibco), 10 ng/ml BDNF (Thermo Fisher Scientific, Cat# 17834733), 10 ng/ml NT-3 (Thermo Fisher Scientific, Cat# 17894523), and 1 μg/ml laminin (Roche). iNeurons were cultured until DIV24 with weekly half-medium changes.

### HEK293T cell culture, lentivirus packaging and lentiviral transduction

HEK293T cells were maintained in Dulbecco’s Modified Eagles Medium (DMEM) high glucose with stable glutamine and sodium pyruvate (Capricorn Scientific) supplemented with 10% fetal bovine serum (GIBCO) and 1% Penicillin-Streptomycin (GIBCO). The cells were maintained at 37°C in 5% CO2.

For mass spectrometry experiments with V5-APEX2-SEC13, lentiviruses were produced by transient transfection of HEK293T cells with transfer vector containing V5-APEX2-SEC13, packaging vector psPAX2 and envelope vector pMD2.G with a ratio of 1:1:1 using PEI Max (Polysciences) according to standard protocols with a 3:1 PEI Max:DNA ratio. PEI Max/DNA complexes were prepared in serum-free Opti-MEM (GIBCO) and added to the cells after 20 min. Medium was completely removed after 4 h and changed to NB supplemented with 0.5 mM glutamine. Supernatants from packaging cells were collected 48-72 h post-transfection, filtered through a 0.45 μm filter, concentrated with Ultra Centrifugal Filter 100 kDa MWCO (AMICON) at 4°C 1000g for 45 min. Concentrated lentivirus was resuspended in appropriate volumes of fresh NB medium and immediately used to transduce DIV4 cortical neurons grown on 6-well plates in NB medium supplemented with 2% B27 (Gibco), 0.5 mM Glutamine (Gibco), 15.6 μM Glutamate (Sigma), and 1% Penicillin/Streptomycin (Gibco). Cells were maintained at 37 °C in 5% CO2 until further processing at DIV9.

For shRNA validation, lentiviruses were produced by transient transfection of HEK293T cells with transfer vector pLKO.1 puro containing shRNA, packaging vector psPAX2 and envelope vector pMD2.G following same protocol as above.

### DNA and shRNA Constructs

The following plasmids were used: FUGW was a gift from David Baltimore (Addgene plasmid #14883)^89^, pLKO.1 puro was a gift from Bob Weinberg (Addgene plasmid #8453)^90^, Str-KDEL_SBP-EGFP-Ecadherin was a gift from Franck Perez (Addgene plasmid #65286)^33^, psPAX2 and pMD2.G were gifts from Didier Trono (Addgene plasmids #12260 and #12259), pmScarlet-i-C1 was a gift from Dorus Gadella (Addgene plasmid #85044)^91^, Halo-TfR-RUSH and SAR1B-H79G-mOrange2 were gifts from Jennifer Lippincott-Schwartz (Addgene plasmid #166905 and #166900)^79^, PB-Ef1a-PP7 coat protein-Halo-GB1x3-WPRE was a gift from Michael Ward (Addgene plasmid # 198337)^24^, pPB-cytERM-SUNTAG-MS2 was a gift from Jennifer Lippincott-Schwartz (Addgene plasmid # 247298)^55^, GFP-MAPPER was a gift from Jen Liou (Addgene plasmid # 117721)^58^, pEGFP-SEC22B was a gift from Thierry Galli (Addgene plasmid #101918)^57^, pcDNA3.1-MS2-HA-HsNBAS_AA was a gift from Elisa Izaurralde (Addgene plasmid #148236), mito-V5-APEX2 was a gift from Alice Ting (Addgene plasmid #72480)^46^, pEGFP-SEC23A and pEYFP-SEC24D were gifts from David Stephens (Addgene plasmid #66609 and #66614)^92^, pDEST47-SAR1-GFP was a gift from Richard Kahn (Addgene plasmid # 67409)^93^, pORANGE Cloning template vector was a gift from Harold MacGillavry (Addgene plasmid #131471)^94^, H2B-mNeonGreen-IRESpuro2 was a gift from Daniel Gerlich (Addgene plasmid #183745)^95^, pEGFP(A206K)-C1 and pEGFP(A206K)-N1 was a gift from Dr. Jennifer Lippincott-Schwartz, GW1-GFP-ZW10 was a gift from Peter van der Sluijs^96^, HA-NF186-mRFP-FKBP was a gift from Casper Hoogenraad^97^, HA-BicD2-FRB was a gift from Lukas Kapitein^43^, Halo-CLCa was a gift from Harold MacGillavry^98^, pmCherry-N1 and mCherry-C1 (Clontech), pGW2-mCherry and pGW1-BFP were gifts from Dr. Lukas Kapitein^43^. The following shRNA inserted in pSuper vector was used in this study: VAPA shRNA #1 (5’-GCATGCAGAGTGCTGTTTC-3’)^99^.

The following plasmids have been previously described: V5-SEC16B and Halo-SEC61β^100^, HA-KIFC1-Strep, mCherry-SBP-RTN4, P180-ΔCoiled-coil-GFP^37^

The plasmids generated in this study include:

For RUSH-KDEL-SYT1-mNeonGreen, the STIM1sp-Strep-KDEL-intron-IRES fragment was PCR amplified from Str-KDEL_SBP-EGFP-Ecadherin (Addgene plasmid # 65286) and inserted in the FUGW vector (Addgene plasmid # 14883) between XbaI and EcoRI restriction sites using Gibson Assembly (New England Biolabs). This created an FUGW-Strep-KDEL-intron-IRES (named FUGW-RUSH). FUGW-RUSH was further digested with EcoRI and BamHI. The SBP-HA fragment was PCR amplified from an SBP-HA-APPwt-mNG construct (a gift from D. Draper, Farías Lab); SYT1 was PCR amplified from rat cortical neuron cDNA; and the mNeonGreen fragment was amplified from the mNeonGreen-N1 vector (Farías lab), flanked by AgeI and SmiI restriction sites. These fragments were further inserted in digested FUGW-RUSH vector by Gibson assembly and flexible linkers were added between SBP-HA and SYT1, as well as between SYT1 and mNeonGreen. The following primers were used to generate this construct:

FUGW-XbaI Fw: 5 - ttgggctgcaggtcgactctagagccaccatggatgtatgcgtccgtc - 3’

FUGW-EcoRI Rv: 5’ - gctagcttcgaagaattcttatcatcgtgtttttcaaaggaaaacca - 3’

SBP-HA Fw: 5’ - cctttgaaaaacacgatgataaggccaccatgggtagcgacgagaagaccactgg - 3’

SBP-HA Rv: 5’ - gctaccgctgccgctaccgc - 3’

SYT1 cDNA Fw: 5’ - gtagcggtagcggcagcggtagcatggtgagtgccagtcatcc - 3’

SYT1 cDNA Rv: 5’ - gaattcgccagaaccagcagcggagccagcggatcccttcttgacagccagcatgg - 3’

mNeonGreen Fw: 5’ - gctgctggttctggcgaattcaccggtatggtgagcaagggcgaggaggataa - 3’

mNeonGreen Rv: 5’ - agcttgatatcgaattgttaacgatttaaatttacttgtacagctcgtccatgcc - 3’

To generate RUSH-SYT1-Halo construct, RUSH-SYT1-mNeonGreen was digested with AgeI and SmiI, and mNeonGreen was replaced by a HaloTag fragment, which was PCR amplified from a HaloTag-C1 construct. The following primers were used for HaloTag amplification:

HaloTag Fw: 5’ - cgctgctggttctggcgaattcaccggtatggcagaaatcggtactggctttcca - 3’

HaloTag Rv: 5’ - tgatatcgaattgttaacgatttaaatttagccggaaatctcgagcgtggacag - 3’

For RUSH-P180ΔCC-HA-SYT1-mNG, the STIM1sp-Strep fragment was PCR amplified from Str-KDEL-SBP-eGFP-Ecadherin (Addgene plasmid #65286). Next, the P180ΔCC fragment was PCR amplified from a P180ΔCC-GFP.^37^ The intron-IRES-SBP-HA-SYT1-mNeonGreen fragment was PCR amplified from RUSH-SYT1-mNeonGreen (described above). These fragments were subsequently inserted in the FUGW vector (Addgene plasmid # 14883), between XbaI and EcoRI restriction sites using Gibson assembly. The Kozak sequence was added at the start of the STIM1-sp-Strep fragment and flexible linkers were added between these fragments. The following primers were used to generate this construct:

STIM1sp-Strep Fw: 5’ - gcttgggctgcaggtcgact - 3’

STIM1sp-Strep Rv: 5’ - ggtgctcgagatctgagtccgctgctggacggcatccagag - 3’

P180ΔCC Fw: 5’ - cggactcagatctcgagcaccatggatatttacgacactcaaac - 3’

P180ΔCC Rv: 5’ - cagttatctatgcggccgctcaagaagctgactctgtcttttt - 3’

Intron-SBP-HA-SYT1-mNG Fw: 5’ - tgagcggccgcatagataac - 3’

Intron-SBP-HA-SYT1-mNG Rv: 5’ - acggatccgctagcttcgaagaattcatttaaatttacttgtacagctcgtcc-3’

For RUSH-NF186-mNeonGreen, NF186 signal peptide and NF186 were PCR amplified from HA-NF186-mRFP-FKBP;^97^ SBP-HA was PCR amplified from SBP-HA-APP-mNG construct (a gift from D. Draper, Farías Lab); and mNeonGreen was amplified from the mNeonGreen-N1 vector (Farías Lab) with added AgeI and SmiI restriction sites. These fragments were subsequently inserted in FUGW-RUSH vector between EcoRI and BamHI restriction sites by Gibson assembly. Flexible linkers were added between SBP-HA and NF186, as well as between NF186 and mNeonGreen. The following primers were used to generate this construct:

Kozak-NF186sp Fw: 5’ - cctttgaaaaacacgatgataaggccaccatgatggccaggcagcaggcgcc - 3’

Kozak-NF186sp Rv: 5’ - ggcccctccgaggctgaggag - 3’

SBP-HA Fw: 5’ - tcctcctcagcctcggaggggccggtagcgacgagaagaccactgg - 3’

SBP-HA Rv: 5’ - gctaccgctgccgctaccgc - 3’

NF186 Fw: 5’ - gtagcggtagcggcagcggtagcattgagattccgatggattacc - 3’

NF186 Rv: 5’ -actgcctcgcccttgctcaccataccggtgaattcgccagaaccagcagcggagccagcggatccgg ccagggaatagatggcattgactg - 3’

mNeonGreen Fw: 5’ - gctgctggttctggcgaattcaccggtatggtgagcaagggcgaggaggataa - 3’

mNeonGreen Rv: 5’ - agcttgatatcgaattgttaacgatttaaatttacttgtacagctcgtccatgcc - 3’

For RUSH-Ii-L1CAM-mNeonGreen, we first generated the FUGW-RUSH-Ii vector. FUGW vector (Addgene plasmid # 14883) was digested with BamHI and EcoRI. Strep-Ii-Intron-IRES was amplified from pIRES-neo3-Strep-Ii-LAMP2-SBP-FKBP-GFP (a gift from Judith Klumperman). The resulting PCR product was assembled with digested FUGW using Gibson Assembly to generate FUGW-RUSH-Ii for subsequent cloning purposes. The following primers were used:

Strep-Ii-IRES Fw: 5’ - ttgggctgcaggtcgactctagagccaccatgcacaggaggagaagcag - 3’

Strep-Ii-IRES Rv: 5’ - gctagcttcgaagaattcttatcatcgtgtttttcaaaggaaaacca - 3’

Next, FUGW-RUSH-Ii was digested with EcoRI and BamHI. L1CAM signal peptide and L1CAM was amplified from GW1-PAGFP-mL1CAM (gift from Casper Hoogenraad). An HA tag with a linker was added between the signal peptide and L1CAM sequence, then mNeonGreen was PCR amplified from H2B-mNeonGreen-IRESpuro2 (Addgene plasmid # 183745) and SBP was PCR amplified from Str-KDEL_SBP_EGFP-Ecadherin (Addgene plasmid # 65286). All fragments were inserted into FUGW-RUSH-Ii between EcoRI and BamHI by Gibson Assembly. The following primers were used to generate this construct:

L1CAM sp Fw: 5’ - aaaacacgatgataagaattatggtcgtgatgctgcggta - 3’

L1CAM sp Rv: 5’ - ggaacatcgtatgggtagctaccgagcaggcaggggctgca - 3’

HA-Linker-L1CAM Fw: 5’ - agattacgctggtagcggtagcggcagcatacagattccagacgaatataaaggac-3’

HA-Linker-L1CAM Rv: 5’ - gcagcagatccagcggatccttctagggctactgcaggattg - 3’

mNeonGreen Fw: 5’ - ctggctccgctgctggttctggcgaattcgtgagcaagggcgaggagga - 3’

mNeonGreen Rv: 5’ - cttgtacagctcgtccatgcc - 3’

SBP Fw: 5’ - ggcatggacgagctgtacaagggatccgacgagaagaccactggttgg - 3’

SBP Rv: 5’ - gaattgttaacggatcgaattcttatggttcacgttgaccttgtgg - 3’

For generation of GFP-α-COP: PCR-amplified α-COP fragment from rat cDNA library was inserted into pEGFP-C1 at XhoI and XmaI restriction sites. The following primers were used:

α-COP Fw: 5’-tatctcgagggatgctaaccaaattcgagacc - 3’

α-COP Rv: 5’-tatcccgggttagcgaaactgaagaggactgatc - 3’

For generation of GFP-β-COP, PCR-amplified β -COP fragment from rat cDNA library was inserted into pEGFP-C1 at BspEI and XhoI restriction sites. The following primers were used:

β-COP Fw: 5’ - tattccggaaccatgaccgcagctgagaacgtg - 3’

β-COP Rv: 5’ - tatctcgagttagagactagtcttcttttgag - 3’

For PB-EF1α-SYT1-UTR-PP7 and PB-EF1α-L1CAM-UTR-PP7, the PB-EF1α-ACTB-UTR-PP7 plasmid (a gift from Michael Ward) was digested with NheI and BsrGI. SYT1 was PCR amplified from RUSH-KDEL-SYT1-mNeonGreen (described above) and SYT1 3’ UTR was PCR amplified from rat cDNA. L1CAM was PCR amplified from L1CAM-DHFR-mNeonGreen and L1CAM 3’UTR was PCR amplified from rat cDNA. These fragments were inserted separately into PB-EF1α-ACTB-UTR-PP7 using Gibson Assembly. The following primers were used to generate these constructs:

SYT1 CDS Fw: 5’ - tccatttcaggtgtcgtgagctagcgccaccatggtgagtgccagtcatcc - 3’

SYT1 CDS Rv: 5’ - cagaaaggcttcgttttccctctagctagcttacttcttgacagccagcatgg - 3’

SYT1 3’UTR Fw: 5’ - agggaaaacgaagcctttctg - 3’

SYT1 3’UTR Rv: 5’ - taccttaggatccggtacctgtacaagcatactggtatatacaacagtgt - 3’

L1CAM CDS Fw: 5’ - ttccatttcaggtgtcgtgagctagcgccaccatggtcgtgatgctg - 3’

L1CAM CDS Rv: 5’ - gcctcacatgactggaccttgctagctagcttattctagggctactgcaggattgat - 3’

L1CAM 3’UTR Fw: 5’ - caaggtccagtcatgtgaggc - 3’

L1CAM 3’UTR Rv: 5’ - gtaccttaggatccggtacctgtacattctctggtgggtatcgaagg - 3’

For V5-APEX2-SEC13-mNG, we first generated vectors FUGW-IRES-mNeonGreen and V5-APEX-C1.

For generation of FUGW-IRES-mNG, FUGW (Addgene plasmid # 14883) was first digested with BamHI and EcoRI. Intron-IRES was amplified from Str-KDEL_SBP_EGFP-Ecadherin (Addgene plasmid # 65286), mNeonGreen was amplified from H2B-mNeonGreen-IRESpuro2 (Addgene plasmid # 183745). All fragments were assembled by Gibson assembly into FUGW vector between BamHI and EcoRI with the EcoRI site destroyed. The following primers were used to generate this construct:

Intron-IRES Fw: 5’ - caggtcgactctagaggatcggatccgaattaattcgctgtctgcgag - 3’

Intron-IRES Rv: 5’ - gcccttgctcaccatggttatcatcgtgtttttcaaaggaaaaccacg - 3’

mNeonGreen Fw: 5’ - acgatgataaccatggtgagcaagggcgag - 3’

mNeonGreen Rv: 5’ - tatcgaattgttaacggatcttacttgtacagctcgtccatgcc - 3’

For generation of V5-APEX2-C1 vector, V5-APEX2 was PCR amplified from mito-V5-APEX2 (Addgene plasmid #72480) and inserted into pEGFP(A206K)-C1 vector between BglII and AgeI restriction sites using the following primers:

V5-APEX2-linker Fw: 5’ - cgtcagatccgctagcgctaccggtGCCACCATGGGTAAGCCTATC - 3’

V5-APEX2-linker Rv: 5’ - aattcgaagcttgagctcgagatctGCTACCGCTGCCGCTACC - 3’

Then, SEC13 sequence was PCR amplified from rat cDNA, and V5-APEX2 with a Kozak sequence in front, was amplified from pV5-APEX2-C1. These amplified sequences were subsequently inserted in FUGW-IRES-mNeoGreen between XbaI and BamHI restriction sites using Gibson assembly. A flexible linker was added in front of the SEC13 sequence. The following primers were used to generate FUGW-V5-APEX2-SEC13:

Kozak-V5-APEX2 Fw: 5’- gcttgggctgcaggtcgactgccaccatgggtaagcctatc - 3’

Kozak-V5-APEX2 Rv: 5’ - gccgctgccgctggtgctgccggatcccttggcatcagcaaacccaag - 3’

SEC13 Fw: 5’ - aagggatccggcagcaccagcggcagcggcatggtgtcagtaattaacac - 3’

SEC13 Rv: 5’- gcagacagcgaattaattcggaattccgtcattgctcattctgttg - 3’

For generation of GFP-SEC16A knock-in, 20 nt RNA guides were designed and selected using the UCSC genome browser gateway (https://genome-euro.ucsc.edu/), with RGSC5.0/rn5 (Rattus norvegicus) as reference genome and NGG (SpCas9, 3’ side) PAM. Target sequences were chosen, taking into consideration the MIT guide specificity score.^101^ The chosen targets were inserted in the pORANGE template backbone^94^ at BbsI restriction sites. The donor sequence was designed to contain a GFP tag flanked by two Cas9 target sites and PCR amplified from a GFP containing plasmid. To facilitate genomic integration of the donor sequence in the correct orientation, these target sites including PAM sequences were inserted as the reverse complement of the genomic target sequence. This donor sequence was then ligated in the pORANGE vector containing targets at HindIII and XhoI restriction sites. Short linker sequences of at least three amino acids and additional base pairs to make the donor in frame after integration in the genome were introduced between the target sites and the tag sequence. The primers for SEC16A target site and donor were as followed:

SEC16A target Fw: 5’ - aaacttataagcagcatctcctgc - 3’

SEC16A target Rv; 5’ - caccgcaggagatgctgcttataa - 3’

GFP donor Fw: 5’-ataaagcttgcaggagatgctgcttataaaggcagccaccatggctagcggagtgagcaagggcgaggag-3’

GFP donor Rv: 5’-atactcgagcctttataagcagcatctcctgcagagcgcttccactcttgtacagctcgtccatgc-3’

For V5-SEC31A, we first generated pCMV-V5-C1 vector by inserting a V5 fragment into pCMV-mScarletii-C1 vector (Addgene plasmid #85044) between AgeI and BsrGI restriction sites. SEC31A fragment was PCR amplified from pEGFP-SEC31A (Addgene plasmid #66612) and inserted into pCMV-V5-C1 vector generated above between BspEI and BamHI restriction sites. A flexible linker (GPKLGSGS) was also included in front of SEC31A sequence. The following primers were used to generated pCMV-V5-C1 and V5-SEC31A:

Kozak-V5-C1 Fw: 5’ - cgtcagatccgctagcgctaactgccaccatgggtaag - 3’

Kozak-V5-C1 Rv: 5’ - cgagatctgagtccggacttgggccccgtagaatcgag - 3’

SEC31A Fw: 5’ - cgattctacggggcccaagttaggcagcggtagtatgaagttaaaggaagtagatc - 3’

SEC31A Rv: 5’ - tcagttatctagatccggtgttagacacccagcttattg - 3’

For Halo-SEC24D, we first generated a pHaloTag-C1 vector. Briefly, HaloTag was PCR amplified from Halo-CLCa (a gift from Harold MacGillavry), and inserted in pGFP(A206K)-C1 vector digested with AgeI and BglII restriction enzymes using Gibson Assembly. The following primers were used for amplification of HaloTag:

Halo Fw: 5’ - cgtcagatccgctagcgctagccaccatggcagaaatcggtactggc - 3’

Halo Rv: 5’ - aattcgaagcttgagctcgagatctgctaccgctgccgctacc - 3’

HaloTag was then PCR amplified from HaloTag-C1 vector and SEC24D was amplified from pEYFP-SEC24D (Addgene plasmid #66614). These fragments were inserted into the pEGFP(A206K)-C1 between AgeI and EcoRI restriction sites. A kozak sequence and a flexible linker were inserted by addition to cloning primers. The following primers were used to generate this construct:

HaloTag Fw: 5’ - gatccgctagcgctaccggtcgccaccatggcagaaatcggtactggcttt - 3’

HaloTag Rv: 5’ - agaaccagcagcggagccagcggatccgccggaaatctcgagcgtgga - 3’

SEC24D Fw: 5’ - cgctggctccgctgctggttctggcgaattcatgagtcaacaaggttacgtggct - 3’

SEC24D Rv: 5’ - attcgaagcttgagctcgagagcggatccttaattaagcagctgacagatctcc - 3’

For SAR1B-H79G-GFP, SAR1B-H79G was PCR amplified from SAR1B-H79G-mOrange2 (Addgene plasmid #166900) and inserted into pEGFP(A206K)-N1 between BamHI and BglII restriction sites. A kozak sequence was inserted by addition to cloning primers. The following primers were used to generate this construct:

Sar1B-H79G Fw: 5’ - cgctagcgctaccggactcagatctgccaccatgtccttcatattt - 3’

Sar1B-H79G Rv: 5’ - ccatggtggcgaccggtggatccgaaccgctgccgctaccatcgatgtactgtgccatc - 3’

For the Strep-SBP heterodimerization system, mCh-SBP-SEC23A or GFP-SBP-SEC23A were generated. The SEC23A fragment was PCR amplified from EGFP-SEC23A (Addgene plasmid #66609) and inserted into either mCh-SBP-C1 or GFP(A206K)-SBP-C1 plasmids^37^ between SalI and BamHI restriction sites. The following primers were used:

SEC23A Fw: 5’ - caaagcttcgaattctgcagtcgaccggtagcggcagcggtagcatgacaacctatttggaattcattc - 3’

SEC23A Rv: 5’ - tcagttatctagatccggtggatcctcaagcagcactggacac - 3’

For the FKBP-FRB heterodimerization system, FLAG-FRB-KIFC1 and mCh-FKBP-SEC23A or GFP-FKBP-SEC23A were generated.

For FLAG-FRB-KIFC1, FUGW vector (Addgene plasmid #14883) was digested with XbaI and EcoRI. KIFC1 was first subcloned from Strep-KIFC1-HA^37^, FRB was subcloned from HA-BicD2-FRB (a gift from Lukas Kapitein)^43^, a 20 amino acid linker GSAGSAAGSGAGSAAGSGEF was introduced behind the FRB fragment. An additional FLAG tag was introduced in front of FRB-KIFC1 using primers. All fragments were inserted into restricted FUGW vector by Gibson assembly. These primers were used to generate this construct:

KIFC1 Fw: 5’ - tacaaagacgatgacgacaagaccggtatcctctggcatgagatgtggcatgaag - 3’

KIFC1 Rv: 5’ - cgctagcttcgaagaattcttacttcctgttggcctgagcagt - 3’

FRB Fw: 5’ - tacaaagacgatgacgacaagaccggtatcctctggcatgagatgtggcatgaag - 3’

FRB Rv: 5’ - gcagcggagccagcggatccctttgagattcgtcggaacacatgataatagagg - 3’

20aa linker: 5’ - gaattcgccagaaccagcagcagatccagcgccagaacctgcagcggagccagcggatcc - 3’

FLAG tag: 5’ - gctgcaggtcgactctagagccaccatggactacaaagacgatgacgacaagaccggt - 3’

For mCh-FKBP-SEC23A or GFP-FKBP-SEC23A, SEC23A was PCR amplified from EGFP-SEC23A (Addgene plasmid #66609) and 2x FKBP was amplified from mCh-FKBP2x-RTN4^37^. These fragments were inserted into either mCherry-C1 or GFP(A206K)-C1 vector, between SalI and BglII restriction sites. A linker was added between FKBP and SEC23A. The following primers were used to generate these constructs:

FKBP2x Fw: 5’ - gctgtacaagtccggactcagatctggagtgcaggtggaaaccatc - 3’

FKBP2x Rv: 5’ - aggttgtcatgctaccgctgccgctacc - 3’

SEC23A Fw: 5’ - cagcggtagcatgacaacctatttggaattcattc - 3’

SEC23A Rv: 5’ - atcccgggcccgcggtaccgtcgactcaagcagcactggacac - 3’

For Halo-NBAS and Halo-ZW10, NBAS was PCR amplified from pcDNA3.1-MS2-HA-HsNBAS_AA (Addgene plasmid # 148236), and ZW10 was PCR amplified from GFP-GW1-ZW10^96^. These fragments were inserted separately in HaloTag-C1 vector (described above) digested with BglII and BamHI. The following primers were used to generate these two plasmids:

NBAS Fw: 5’ - cggtagcggcagcggtagcagatctatggcggcccccgagtca - 3’

NBAS Rv: 5’ - agttatctagatccggtggatcctcacacccagtgctgtgctgc - 3’

ZW10 Fw: 5’ - cggtagcggcagcggtagcagatctatggcctcgttcgtgaca - 3’

ZW10 Rv: 5’ - cggtagcggcagcggtagcagatctatggcctcgttcgtgaca - 3’

For generation of ALFA-SEC22B (1-110), ALFA was PCR amplified from Xph20-ALFA (a gift from Harold MacGillavry). SEC22B (1-110) fragment was PCR amplified from GFP-SEC22B^57^. These fragments were inserted in GFP-C1 digested with AgeI and EcoRI. The following primers were used to generate this plasmid:

ALFA Fw: 5’ - tcagatccgctagcgctaccggtgccaccatgggctctcgtcttgaagaggaacttc - 3’

ALFA Rv: 5’ - gccagaaccagcagcggagccagcggatccttcagtaagacgtctacgaag - 3’

SEC22B (1-110) Fw: 5’ - gctgctggttctggcgaattcgtgctgctgacaatgattgcc - 3’

SEC22B (1-110) Rv: 5’ - ccgcggtaccgtcgactgcagaattctcaatagggcctagacacagtggg - 3’

The following rat shRNA-targeting sequences inserted to pLKO.1-puro were used in this study:

scramble shRNA (5’-gatgaaatattccgcaagtaa-3’) or

scramble shRNA (5’-gaaacaaatcggaaatctt-3’)

HDLBP-shRNA (5’-atggtcaaagacctgattaat-3’)

SEC22B-shRNA (5’-tcatcatgctgatagtgtatg-3’)

ZW10-shRNA (5’-gactctgtcgtcctaaatttg-3’)

NBAS-shRNA (5’-atgaccagaaaggccattaag-3’)

The following sequences were used in this study to validate HDLBP, SEC22B, ZW10 and NBAS shRNA-mediated knockdown efficiency:

For HDLBP:

HDLBP qPCR Fw1: 5’ - agagaggaaaacccagttcaca - 3’

HDLBP qPCR Rv1: 5’ - tcggtctccttggcctctc - 3’

For SEC22B:

SEC22B qPCR Fw2: 5’ - agaagcactctcagcattgga - 3’

SEC22B qPCR Rv2: 5’ - cttcgcatcctggcggtatt - 3’

For ZW10:

ZW10 qPCR Fw1: 5’ - aaatcggagatcggtgggc - 3’

ZW10 qPCR Rv1: 5’ - gcagccgctcttctttctgt - 3’

For NBAS:

NBAS qPCR Fw2: 5’ - ccagtcaccatgaggtcgag - 3’

NBAS qPCR Rv2: 5’ - acaggttgttggctatgacaca - 3’

Primers targeting GAPDH were used as internal control.

GAPDH qPCR Fw: 5’ - caactccctcaagattgtcagcaa - 3’

GAPDH qPCR Rv: 5’ - ggcatggactgtggtcatga - 3’

### Antibodies and reagents

The following primary antibodies were used in this study: mouse anti-Puromycin (Merck Cat# MABE343, RRID: AB_2566826, 1:1000), mouse anti-SMI-312 (BioLegend Cat#837904, RRID:AB_2566782, 1:1000), chicken anti-MAP2 (Abcam Cat#ab5392, RRID:AB_2138153, 1:5000), rabbit anti-Synaptotagmin-1 (Synaptic Systems Cat#105 008, RRID:AB_2832236, 1:1000), rabbit anti-Neurexin-1alpha (Frontier Institute/ Nittobo Medical Cat# MSFR104630, 1:300), rabbit anti-L1CAM (Proteintech Cat# 20659-1-AP, RRID:AB_2878717, 1:250), rabbit-anti-TRIM46 (a gift from Casper Hoogenraad,, 1:500), mouse anti-V5 (Thermo Fisher Scientific Cat# R960-25, RRID:AB_2556564, 1:500), rat anti-HA (Roche Cat# 11867423001, RRID:AB_390918, 1:200), mouse anti-GM130 (BD Biosciences Cat#610823, RRID:AB_398142, 1:1000), sheep anti-TGN38 (Bio-Rad Cat#AHP499G, RRID:AB_2203272, 1:1000), mouse anti-ERGIC-53 (Sigma-Aldrich Cat# SAB4200585, 1:500), mouse anti-CALN1 (Abnova Cat# H00083698-A01, RRID:AB_535135, 1:100), guinea pig anti-Synaptophysin1 (Synaptic Systems Cat# 101 004, RRID:AB_1210382, 1:500), mouse anti-GFP (Thermo Fisher Scientific Cat#A-11120; RRID:AB_221568, 1:700), mouse anti-mCherry (Takara Bio, Cat#CL 632543, 1:1000), rabbit anti-SEC31A (Cell Signaling Technology Cat#13466, RRID:AB_2798228, 1:300), mouse anti-β-Tubulin III (Sigma-Aldrich Cat#T8660, RRID:AB_477590, 1:1000), rabbit anti-HDLBP (Proteintech Cat#15406-1-AP, RRID:AB_2117367, 1:500), rabbit anti-NBAS (Proteintech Cat#14683-1-AP, RRID:AB_1959590, 1:200), rabbit anti-ZW10 (Abcam Cat#ab21582, RRID:AB_779030, 1:500), rabbit anti-Synapsin-I (Millipore Cat#AB1543, RRID:AB_2200400, 1:500), mouse anti-RTN4A (named NogoA) (R and D Systems Cat# MAB3098, RRID:AB_10997139), mouse anti-Pan-Neurofascin external (clone A12/18; UC Davis/NIH NeuroMab, Cat#75-172, RRID: AB_2282826, 0.18 mg/ml).

The following secondary antibodies were used in this study: goat anti-mouse Alexa405 (Molecular Probes, Cat#A31553, RRID:AB_221604, 1:500), goat anti-Mouse IgG (H+L) Highly Cross-Adsorbed Alexa Fluor 488 (Thermo Fisher Scientific Cat#A-11029, RRID:AB_2534088, 1:1000), goat anti-mouse Alexa568 (Thermo Fisher; Cat#A-11031, RRID:AB_144696, 1:1000). goat anti-rabbit IgG (H + L) Highly cross-absorbed secondary antibody Alexa Fluor 405 (Thermo Fisher Scientific Cat#A-31556, RRID:AB_221605, 1/500), goat anti-mouse IgG2a cross-absorbed secondary antibody Alexa Fluor 594 (Thermo Fisher Scientific Cat#A-21135, RRID:AB_2535774, 1/1000), goat anti-Mouse IgG (H+L) Highly Cross-Adsorbed Secondary Antibody, Alexa Fluor 647 (Thermo Fisher Scientific Cat#A-21236, RRID:AB_2535805, 1/1000), goat anti-rabbit IgG (H + L) Highly cross-absorbed secondary antibody Alexa Fluor 405 (Thermo Fisher Scientific Cat#A-31556, RRID:AB_221605, 1/500), goat anti-Rabbit IgG (H+L) Highly Cross-Adsorbed Secondary Antibody, Alexa Fluor488 (Thermo Fisher Scientific Cat#A-11034, RRID:AB_2576217, 1/1000), goat anti-rabbit IgG (H+L) Highly Cross-Adsorbed CF®594 (Sigma-Aldrich, Cat#SAB4600110, 1:500), goat anti-rabbit IgG (H + L) highly cross-absorbed secondary antibody Alexa Fluor 568 (Thermo Fisher Scientific Cat#A-11036, RRID:AB_10563566, 1/1000), goat anti-Rabbit IgG (H+L) Highly Cross-Adsorbed Secondary Antibody, Alexa Fluor647 (Thermo Fisher Scientific Cat#A-21245, RRID:AB_2535813, 1:500), goat anti-chicken IgY (H+L) Secondary Antibody, Alexa Fluor488 (Thermo Fisher Scientific Cat#A-11039, RRID:AB_2534096, 1/1000), goat anti-Chicken IgY (H+L) Secondary Antibody, Alexa Fluor568 (Thermo Fisher Scientific Cat#A-11041, RRID:AB_2534098, 1/1000), donkey anti-Sheep IgG (H+L) Cross-Adsorbed Secondary Antibody, Alexa Fluor488 (Thermo Fisher Scientific Cat#A-11015, RRID:AB_2534082, 1/1000), goat anti-Rat IgG (H+L) Cross-Adsorbed Secondary Antibody, Alexa Fluor568 (Thermo Fisher Scientific Cat#A-11077, RRID:AB_2534121, 1/500), Goat anti-Guinea Pig IgG (H+L) Highly Cross-Adsorbed Secondary Antibody, Alexa Fluor 568 (Thermo Fisher Scientific Cat# A-11075, RRID:AB_2534119, 1/1000), Alexa Fluor555 conjugated Strep (Thermo Fisher Scientific Cat#s21381, RRID:AB_2307336, 1:1000), Atto488 conjugated Fluotag-X4® GFP nanobody (NanoTag Biotechnologies, Cat#N0304-At488-L, 1:250), Atto488 conjugated GFP-Booster nanobody (ChromoTek, Cat#gba488, 1:250), Star635P conjugated FluoTag®-X2 anti-ALFA nanobody (NanoTag Biotechnologies, Cat# N1502-Ab635P-L, 1:250), rabbit anti-mouse immunoglobulins/HRP (Agilent Cat# P0260, RRID:AB_2636929, 1/10000), IRDye 800CW Streptavidin (LI-COR Biosciences, Cat#926-32230, 1/10000).

Other reagents used in this study were: Puromycin dihydrochloride (Sigma-Aldrich, Cat#P8833), Anisomycin (Sigma-Aldrich, Cat#9789), Lipofectamine 2000 (Thermo Fisher Scientific, Cat#1639722), Fluoromount-G Mounting Medium (ThermoFisher Scientific, Cat#00-4958-02), anhydrous-DMSO (Thermo Fisher Scientific, Cat#D12345), biotin (Sigma-Aldrich, Cat#B4501), biotin-phenol (Iris Biotech, Cat#LS.3500), H_2_O_2_ (Sigma-Aldrich, Cat#H1009), Sodium Azide (Merck, Cat#K43547688), Sodium L-ascorbate (Sigma, Cat#0000374819), Trolox (Sigma, Cat#238813), rapalog (A/C Heterodimerizer) (Takara Bio, Cat#635057), NeutrAvidin (Thermo Fisher Scientific, Cat#31000), Brefeldin A (Sigma-Aldrich, Cat#B5936), NAD+ (Sigma-Aldrich, Cat# 10127965001), Mix-n-Stain CF640R antibody labeling kit (Biotium, Cat#92245), Duolink® In Situ Detection Reagents FarRed (Sigma-Aldrich, Cat#DUO92013), Polyethylenimine (PEI MAX; Polysciences, Cat#24765), and Triton-X-100 (Sigma, Cat#93433). JFX554 and JFX650 were kindly provided by the Lavis Lab (Janelia)^103^.

### Drug treatments

BFA was used at 2-2.5 μg/ml for 30 min (**Fig. 3n, Supplementary Fig. 3a, 3b, 3f**, and **3g**) or 4h 30 min (**Fig 2c, 2m, 3j, 4h, 5e, 5g**, and **5i**, and **Supplementary Fig. 3c, 3e**, and **5a**) for fixed cell experiments. For live-cell experiments, neurons were incubated with BFA 2-2.5 μg/ml for 30 min and transferred to a spinning disk confocal microscope. Samples were imaged for up to 3h (**Fig. 5c, 5d**, and **Supplementary Fig 6j**). BDNF was used at 100 ng/ml for 30 min, after which cells were incubated with puromycin for 10 min (**Fig. 1d** and **1f**) or kept for 4h with BFA and biotin (**Fig. 2c, 2m, 3j, 4h, 5e, 5g, 5i** and **Supplementary Fig. 5a**) before fixation, or fixed immediately (**Fig. 3c**).

For experiments combining expression of RUSH-SYT1 construct with shRNAs for 4 days, NAD+ (1mM) was added to culture medium to ensure neuron survival in biotin-free medium over long period (**Fig. 5e, 5g** and **Supplementary Fig. 9e**) .

For the Streptavidin/SBP heterodimerization system assay, NeutrAvidin (0.3 mg/ml) was added to neurons immediately after transfection to avoid Strep-SBP uncoupling by the presence of biotin in culture medium. NeutrAvidin was kept in the medium for 48h (**Fig. 3f, 4l, 6a, Supplementary Fig. 7b, 7d**, and **7f**) or 72h (**Fig. 6g**) until fixation.

For drug-induced FKBP/FRB heterodimerization assay, rapalog (500 nM) was added after transfection for 16h before further BFA, biotin and BDNF treatments (**Fig. 3i, 3j**). Ethanol 0.1% was used as a vehicle in control treatment.

### RNA extraction, reverse transcription and quantitative PCR

Rat cortical neurons DIV9 were used to validate knockdown efficiency of HDLBP, ZW10, NBAS, and SEC22B shRNAs. Total RNA from transduced rat cortical neurons was extracted using RNeasy Mini kit (Qiagen, 74104). cDNA was reverse-transcribed using SuperScript III First-Strand Synthesis System (Invitrogen, 18080-051) according to manufacturer’s instructions. Quantification of target genes was done by qPCR using Power SYBR Green PCR Master Mix (Applied Biosystems, 4367659) and ViiA 7 Real-Time PCR System (Applied Biosystems). Data was extracted from ViiaA 7 Real-Time PCR system using QuantStudio Software V1.3 (Applied Biosystems).

### Mass Spectrometry sample preparation and analysis

#### Sample preparation

Neurons transduced with lentivirus at DIV4 stably expressing FUGW-APEX2-SEC13-mNG were subjected to APEX2 treatment at DIV9 as previously described.^104^ Briefly, neurons were incubated with NB media supplemented with 500μM biotin-phenol (IrisBio-tech) at 37°C for 20 min before addition of 1mM H_2_O_2_ at room temperature for 1 min to activate peroxidase activity. Biotinylation was immediately quenched by two washes in ice-cold quencher solution (10mM sodium azide, 10mM sodium ascorbate and 5mM Trolox in HBSS) and incubated 10 min on ice in azide-free quencher solution. Cells were scrapped and pooled according to their respective conditions. Pellets were stored at -80°C until all three biological replicates were collected. Cell pellets were lysed with freshly made quenching-RIPA buffer (150mM NaCl, 50mM Tris-HCl pH7.4, 0.1% SDS, 0.5% sodium-deoxycholate, 1% Triton X-100, 10mM sodium azide, 10mM sodium ascorbate, 5mM Trolox and 1x protease inhibitors (Roche)) on ice for 30 min. Lysates were cleared by centrifuging at 16,000 x g 4°C, supernatants were incubated overnight with pre-equilibrated Pierce magnetic streptavidin beads (Invitrogen) on a rotor at 4°C. Beads were washed three times with detergent-free RIPA buffer (150mM NaCl, 50mM Tris-HCl pH7.4) and three times with freshly prepared 3M Urea buffer (Urea in 50mM ammonium bicarbonate). Beads were resuspended in 10μL of 3M Urea buffer and reduced with 5mM TCEP (Sigma) at room temperature for 30 min, alkylated with 10mM IAA (Sigma) in the dark at room temperature for 20 min and quenched with 20mM DTT (Sigma). Samples were washed three times with 2M Urea buffer and resuspended in 100μL 2M Urea buffer. Suspended beads were first incubated with 1μg LysC for 4 hours at room temperature, followed by adding 1μg Trypsin for digestion at 37°C overnight. Peptides were collected by combining digested supernatant with two subsequent 100μL 2M Urea buffer washes and immediately acidified with 1% trifluoroacetic acid. Digested peptides were desalted on Sep-Pak C18 Cartridges (Waters) and vacuum concentrated for storage until subsequent MS analysis.

#### Western blot analysis

Cell lysates were prepared as previously described before incubation with beads. 5% of lysates were reserved for immunoblotting. Samples were loaded on a homemade 10% Bis-Acrylamide gels (Bio-Rad) and transferred by wet transfer onto PVDF membranes (Bio-Rad). Membranes were blocked with 5% skimmed milk in TBS-T for 1h at room temperature and were incubated overnight at 4°C with primary antibodies diluted in antibody buffer (3% BSA in TBS-T) on a rotator. Blots were washed 3 times with TBS-T for 5 min each on a shaker and were incubated with anti-mouse HRP-conjugated secondary antibodies in antibody buffer for 45 min at room temperature, then washed three more times with TBS-T. Detection was performed using Clarity Western ECL Substrate (Bio-Rad, Cat#1705060) and imaged on an ImageQuant 800 system (AMERSHAM). To detect biotinylation, membranes were washed 3 times in TBS-T and incubated with IRDye 800CW Streptavidin for 45 min at room temperature. Finally, blots were washed 3 times with TBS-T for 5 min each and twice briefly in TBS before developing on an Odyssey CLx imaging system (LICOR) with Image Studio version 5.2.

#### MS and data analysis

All samples were reconstituted in 0.1% formic acid and analyzed on a Orbitrap Exploris 480 mass spectrometer (Thermo Fisher Scientific, San Jose, CA, United States) coupled to an UltiMate 3000 UHPLC system (Thermo Fisher Scientific, San Jose, CA, United States). Peptides were loaded onto a trap column (C18 PepMap100, 5 μm, 100 Å, 5 mm × 300 μm, Thermo Fisher Scientific, San Jose, CA, United States) with solvent A (0.1% formic acid in water) at 30 μl/min flowrate and chromatographically separated over the analytical column (Poroshell 120 EC C18, Agilent Technologies, 50 μm x 75 cm, 2.7 μm) using 180 min gradient at 300 nL/min flow rate. The gradient proceeds as follows: 9% solvent B (0.1% FA in 80% acetonitrile, 20% water) for 1 min, 9−13% for 1 min, 13−44% for 155 min, 44−55% for 5 min, 55−99% for 5 min, 99% for 3 min, and finally the system equilibrated with 9% B for 10 min.

The mass spectrometers were used in a data-dependent mode, which automatically switched between MS and MS/MS. After a survey MS scan ranging from 375 to 1600 m/z with 14 s dynamic exclusion time, the most abundant peptides of 120 m/z or higher were subjected to high energy collision dissociation (HCD) for further fragmentation using a 1.4 m/z isolation window. MS spectra were acquired in high-resolution mode (R > 60,000), whereas MS2 was in high-sensitivity mode (R > 15,000) and 28% normalized collision energy.

For data analysis, raw files were processed using MaxQuant’s (Version 2.0.1.0).^105^ Andromeda search engine in reversed decoy mode based on rat reference proteome (Uniprot-FASTA, UP000002494, downloaded March 2023) with an FDR of 0.01 at both peptide and protein levels. Digestion parameters were set to specific digestion with trypsin with a maximum number of 2 missed cleavage sites and a minimum peptide length of 7. Oxidation of methionine and amino-terminal acetylation were set as variable and carbamidomethylation of cysteine as fixed modifications. The tolerance window was kept at default. A total of 3 biological replicates with 2 technical replicates each were analyzed. The resulting protein group file was first processed using Perseus (version 1.6.15.0)^106^ to filter for common contaminants, reverse site-specific identifications. Proteins with peptide count <2 were excluded. The processed file was migrated to R (Version 4.5.0) for further analysis with a custom script. Briefly, raw intensity was log_2_ transformed and only proteins found in at least four replicates in the same condition were retained for analysis. Control and H_2_O_2_ conditions were separated and missing values within the same condition were imputed using BPCA and median normalized. Next, both conditions were joined back. For data visualization, proteins only found in H_2_O_2_ but not in control were imputed using QRILC for statistical purposes based on the assumption that these proteins were biologically relevant to SEC13 interactome. APEX2-SEC13 samples were tested using unpaired two sample *t*-test and proteins with a p value < 0.05 and log_2_ fold change ≥1 were considered significantly enriched. Data were visualized using R (Version 4.5.0).

### Puromycilation assay

To label newly synthesized proteins, neurons were briefly incubated with 10 μM puromycin for 10 min. This duration is commonly used to detect newly made proteins in axons.^17,107^ Following the incubation, any unincorporated puromycin was removed by washing neurons twice with NB medium. Neurons were then immediately fixed and processed for immunostaining as detailed below. For control conditions, neurons were pre-treated with 100 μM anisomycin for 40 min.

### Puromycilation-proximity ligation assay (Puro-PLA)

Newly synthesized proteins by proximity ligation were detected using anti-puromycin antibody in combination with antibodies recognizing specific POIs. Detection was performed using Duolink® PLA detection reagents (Sigma-Aldrich) according to the manufacturer’s protocol. Duolink® PLA probes, including Duolink® In Situ PLA probe anti-mouse PLUS (Sigma-Aldrich, Cat#DUO92001) and anti-rabbit MINUS (Sigma-Aldrich, Cat#DUO92005) were used as secondary antibodies. Duolink® In Situ Detection Reagents FarRed (Sigma-Aldrich, Cat#DUO92013) were used for ligation, amplification and probe binding.

After puromycilation assay, fixation, permeabilization and washing (as described above), cells were blocked in Duolink® Blocking Solution for 1h at 37°C followed by an incubation with primary antibody pairs diluted in Duolink® Antibody Diluent at 4°C overnight. After washing 2 times, 5 min between each with 1X wash buffer A, PLA probes were added in 1:5 dilution in Duolink® Antibody Diluent for 1h at 37°C. After washing, cells were incubated for 30 min with ligation reaction containing the circularization oligos and T4 ligase diluted in Ligation buffer. After brief washes with wash buffer A, amplification and label probe binding was done with Polymerase and the fluorophore-labeled detection oligo diluted Amplification buffer for 100 min at 37°C. Amplification was stopped by 2 washes in 1X wash buffer B and 1 in 0.01X wash buffer B. For signal stability, cells were washed with PBS/MC fixed for 5 min at room temperature with 4% PFA/sucrose and processed further for immunofluorescence.

### HaloTag labeling

HaloTag labeling was performed with cell-permeable Halo-JFX554 or Halo-JFX650 ligands (Lavis Lab, Janelia)^103^. Ligands were dissolved in DMSO to 50 μM and stored in single-use aliquots at - 20°C. HaloTag ligands were diluted in culture medium to a final concentration of 100-200 nM. Cells were incubated with ligands for 10 min in incubator, at 37°C, 5% CO_2_. After that, cells were rinsed and put back to the culture medium for subsequent fixation (described below) or used for live cell imaging.

### Live HA surface staining

For experiments involving staining for surface HA tag, neurons were incubated with rat anti-HA (Roche) diluted 1:200 in original medium and secondary antibody diluted 1:500 in original medium for 30 min. Incubations were done at 37°C, 5% CO_2_. Cells were then washed in NB buffer and followed subsequent fixation and immunofluorescence protocol.

### Immunofluorescence staining and confocal imaging

Neurons were fixed in 4% paraformaldehyde supplemented with 4% sucrose in PBS for 7-10 min at RT. Fixed cells were washed 3 times with PBS plus magnesium chloride and calcium chloride (PBS-MC) and subsequently permeabilized with 0.2% Triton X-100 in PBS-MC for 15 min at RT. Neurons were again washed with PBS-MC and then incubated with blocking buffer (0.2% porcine gelatin in PBS-MC) for 30 min at 37°C. Next, cells were incubated with primary antibodies diluted in blocking buffer for 30 min at 37°C or overnight at 4°C, washed 3 times with PBS-MC, incubated with secondary antibodies in blocking buffer for 30 min at 37°C, and washed 3 times with PBS-MC. Coverslips were then briefly dipped in Milli-Q water and mounted in Fluoromount-G mounting medium.

Imaging was done using a confocal laser scanning microscope (LSM900, with Zen (blue edition) imaging software version 3.7.97.07000 (Zeiss)) equipped with Plan-Apochromat ×63 NA 1.40 oil DIC and Plan-Apochromat x40 NA1.30 oil DIC objectives. Each confocal image was a z stack of 5−10 images, each averaged 2 times, covering the entire region of interest from top to bottom. Maximum projection images were obtained from the resulting z stack. For fluorescence intensity comparison, settings were kept the same for all conditions.

### Live-cell imaging

Live-cell imaging experiments were conducted using an inverted Nikon Eclipse Ti-E microscope (Nikon) equipped with a Plan Apo VC 100X/NA 1.40 oil objective, an S Fluor 100X/0.5-1.3 oil objective (Nikon) and a Plan Apo VC 60X/NA 1.40 oil objective (Nikon). The system included a Yokogawa CSU-X1-A1 spinning disk confocal unit (Roper Scientific), a Photometrics Evolve 512 EMCCD camera (Roper Scientific), and an incubation chamber (Tokai Hit) mounted on a motorized XYZ stage (Applied Scientific Instrumentation). Coverslips were mounted in a metal ring and imaged using an incubation chamber supplemented with original full medium from neurons. Cells were maintained at optimal conditions of 37°C and 5% CO2 during imaging. Image acquisition and device control were managed using MetaMorph software (version 7.10.2.240). For single-color imaging, each laser channel was exposed for 100−200 ms, while for dual-color imaging, laser channels were sequentially exposed within the same time range. Time-lapse imaging intervals and total durations varied by experiment, as specified in the figure and movie legends. To identify axons, neurons were either co-transfected with a fill and identified by morphology or stained for 15 min with a CF640R-conjugated antibody against the axon initial segment (AIS) protein neurofascin (NF-640R).

### Axon cutting with photoablation laser

Cells were transfected with RUSH-P180ΔCC-HA-SYT1-mNG. Experiments were performed using the ILas2 system (Roper Scientific). Neurons were selected and their positions were saved in MetaMorph software. Teem Photonics 355nm Q-switched pulsed laser was used to perform laser-induced severing. 10% 355nm laser with a spot length of 100 points and diameter of 100 pixels was applied at the proximal axon to separate the axon and the soma. Biotin was immediately added, and images were acquired at distal parts of the severed axons at 1 min intervals for 60-90 min.

### Live-cell imaging with SoRa microscope

For super-resolution imaging with the SoRa microscope, the experiment was performed using a Nikon Eclipse T2 inverted microscope (Nikon) equipped with a CFI SR HP Apo TIRF 100XAC/ NA 1.49 oil objective. The system included a Yokogawa Spinning Disk Field Scanning Confocal Systems CSU-W1 (Nikon) with an added SoRa (Super Resolution by Optical Pixel Reassignment) mode for approximately 1.4x improvement in resolution, an ORCA-Fusion digital sCMOS camera (Hamamatsu), an incubation chamber (Tokai Hit) to maintain samples at 37°C and 5% CO2 during image acquisition. Image acquisition and device control were managed by NIS-Elements (v5.21). Each laser channel was exposed for 400 ms and time-lapse imaging was acquired at 1s intervals for 2 min. Deconvolution on acquired images was performed using the 2D Richardson-Lucy (RL) method with 20 iterations for each channel.

### Image analysis and quantification

#### Puro-PLA signal detection and newly synthesized protein number analysis

To quantify the amount of newly synthesized protein in axons, we acquired multi-channel z-stacks of neurons labelled with puro-PLA signals, endogenous MAP2 for dendrites, and endogenous SMI-312 for axons. Maximum intensity projections were applied to raw images. The SMI-312 signal, excluding the soma plus 20 μm radius from soma edge, was thresholded and this region was added to ROI manager, representing the axonal area. The same region was overlayed on PLA channel, and the ComDet plugin (https://github.com/ekatrukha/ComDet) was used to quantify the number of PLA signals, corresponding to the number of newly synthesized protein. Numbers were normalized to the corresponding area and averaged over multiple cells. Data were presented as number of newly synthesized proteins per 100 μm^2^ axon area in **Fig. 1e, 1g, 1j, 1l** and **1n**.

For knockdown and ERES pulling experiments, the axon was selected as the longest neurite based on a fluorescent protein fill, and a positive signal for the axonal marker SMI-312. A 100-μm axon segment at least 100 μm away from the soma was straightened. The number of newly synthesized proteins was manually quantified (**Fig. 4g** and **4m**).

#### Quantification of RUSH cargo number in axon and percentage of cargo fusing with membrane

To quantify the number of RUSH cargo in the axon, an axon was first identified using the axon initial segment marker TRIM46^102^. A segmented line was traced along the axon, starting at minimum 50 μm away from the soma and straightened for quantifications using Fiji/ImageJ. A 100μm-axon segment was selected, and the number of RUSH cargoes was manually quantified (**Fig. 2d, 3k, 4i, Sf, S1** and **Supplementary Fig. 2e** and **7j**). To assess PM fusion, we calculated the number of RUSH cargoes that were positive for HA surface staining (**Fig. 5k**) and their percentage relative to total cargoes (**Fig. 2n,5h** and **5j**).

For experiments involving axon cutting, axons were identified by the enrichment of P180ΔCC. Videos were acquired at regions of at least 50 μm away from the soma for 1h, at 1 frame every minute. Cumulated number of cargoes were tracked frame-by-frame, new cargoes were defined as puncta absent from all preceding frames, thus excluding pre-existing cargoes (**Fig. 2k**).

#### Quantification of ERES number in axon

The axon was identified based on morphology, using a fill or staining for β-Tubulin or TRIM46. A 100-μm axon segment was analyzed in which number of ERES were manually counted (**Fig. 3d, 3g, 4k, Sn, Sp**, and **Supplementary Fig. 6f**) or by using ImageJ’s build-in Find Maxima function (**Supplementary Fig. 7c** and **7e**). To analyze the colocalization between SEC23A and SEC24D, SEC31A or SEC16B, a 100-μm axon segment was selected for quantification. The ImageJ ComDet plugin (https://github.com/ekatrukha/ComDet) was used on these segments to detect number of puncta and percentage of colocalization between channels (**Fig. 3b**).

#### Kymograph analysis

Kymographs were generated from live cell images using Fiji/ImageJ. Segmented lines were drawn and straightened along the axon identified by morphology using a fill and/or using the AIS marker NF-CF640R. Straightened axons were re-sliced followed by z-projection to obtain kymographs. A segment of 20-30 μm was cropped out for analysis. Anterograde movements were oriented from left to right in all kymographs. Time of recording and length of segments are indicated in each kymograph (**Fig. 4a, Supplementary Fig. 6h**, and **6k**). The number of events for anterograde, retrograde and stationary GFP-SEC23A puncta was manually counted from kymographs (**Supplementary Fig. 6i** and **6l**)

#### Fluorescence intensity profile plots

The distribution of different proteins was analyzed using ImageJ. Plot profiles were generated using a line traced along specified markers. Length of traced profile line is indicated in each intensity plot. Representative images and their corresponding plots are shown in each figure (**Fig. 3p, 3r, 3t, 3v, 6h**, and **Supplementary Fig. 5b** and **5c**).

#### SEC23A and SYT1 mRNA proximity quantification

To quantify the fraction of SEC23A puncta in proximity to SYT1 mRNA (and vice-versa), 3-min videos were acquired from neurons co-expressing SEC23A and SYT1 mRNA reporters. SEC23A and SYT1 mRNA were manually counted frame-by-frame and proximity was defined as close apposition of less than 500 nm. The percentage of SEC23A puncta adjacent to SYT1 mRNA was calculated relative to total SEC23A puncta, reciprocally, the percentage of SYT1 mRNA puncta close to SEC23A was determined relative to total SYT1 mRNA puncta (**Fig. 4b**).

#### ER intensity analysis

To analyze average ER intensity in the axon, a staining for endogenous RTN4A was performed. Samples were imaged with the same settings for laser power, exposure and gain for all conditions. Axons were identified by using a TRIM46 staining. Segmented lines of approximately 40 μm along the axons were manually drawn and straightened. Mean intensity was measured by tracing a line along the straightened axon using Fiji/ImageJ (**Supplementary Fig. 7g**).

#### Enrichment of cargoes in Golgi

To analyze somatic cargo trafficking and sorting to the Golgi, RUSH-SYT1-mNG cargoes were released for 30 min and multi-channel z-stacks were acquired. Maximum-intensity projections were applied to raw images, followed by a staining for endogenous GM130 to mark the Golgi. The GM130 signal was thresholded to create a Golgi mask, which was added to ROI manager, and overlaid onto the RUSH-SYT1-mNG channel. Mean fluorescence intensity within this region was measured. Somatic intensity, excluding the Golgi and background intensity were similarly measured. Golgi enrichment of RUSH-SYT1-mNG cargoes was determined as the ratio of background-subtracted intensities in the Golgi versus the soma (**Supplementary Fig. 9f**).

#### Axon morphology analysis

To analyze axon complexity and morphology, images containing the whole axon were used. Neurons were transfected with a cell fill and the axons were identified using the AIS marker TRIM46. Axon length and morphology (**Fig. 6b, 6c, 6e** and **6f**) were analyzed using NeuronJ plugin for Fiji/ImageJ^108^. Axons were traced and classified as either primary, secondary or tertiary branch. For axons that could not fit in one field of view, several images were taken and stitched together using MosaicJ plugin^109^. All measurements were saved to an Excel file.

#### Synaptic bouton quantification

Presynaptic boutons were identified by swellings along the axon by expressing a cell fill, as described previously^110,111^. Mature boutons were quantified from 3 segments of axons (50 μm each), determined by boutons that were also positive for the endogenous pre-synaptic maker Synapsin-I (**Fig. 6i** and **6k**).

### Statistical analysis

Data processing and statistical analysis were performed using Microsoft Excel, GraphPad Prism (version 10.1.1), MaxQuant (version 2.0.1.0), Perseus (version 1.6.15.0) and R (version 4.5.0). The assumption of data normality was checked using D’Agostino-Pearson omnibus test. Unpaired t-tests, Mann-Whitney tests, Kruskal-Wallis tests followed by Dunn’s multiple comparison test, ordinary one-way ANOVA tests followed by Dunnett’s multiple comparisons or Tukey’s multiple comparisons test were performed for statistical analysis as indicated in figure legends. See Source Data 2 for details on number of experiments, type of analysis and statistical test per experiment.

## Acknowledgements

We thank Professor Dr. Anna Akhmanova (Utrecht University, NL) for providing critical feedback and discussion during the project. This work was supported by the European Research Council (ERC-StG 950617) to G.G.F., Target ALS Foundation (NI-2024-NAI-S5) to G.G.F., Netherlands Organization of Scientific Research (0.16.VIDI.189.019) to G.G.F., and European Research Council (ERC-StG 101163280) to M.K.

## Author contributions

H.H.N. designed and performed experiments, analyzed data, and wrote the manuscript. N.K. designed, cloned constructs and performed experiments to validate the ERES pulling assay. C.H.L. performed mass spectrometry experiments and was supported by M.A. H.J. designed constructs and performed experiments related to axon morphology with heterodimerization system. T.A. and T.L. designed, cloned constructs and performed initial optimizations. D.T.M.N. performed experiments and analyzed qPCR data related to shRNA knockdown. M.B. established human iPSC culture and neuron differentiation protocols and provided iNeurons. M.K. provided feedback and edited the manuscript. G.G.F. designed experiments, supervised the research, coordinated the study, and wrote the manuscript.

## Materials & Correspondence

### Lead contact

Further information and requests for resources, plasmids and reagents should be directed to and will be fulfilled by the lead contact Ginny G. Farías.

### Materials availability

Plasmids in this study will be deposited in Addgene or available upon request as of the date of publication.

### Data and code availability

The mass spectrometry proteomics data generated in this study have been deposited to the ProteomeXchange Consortium via the PRIDE^86^ partner repository with the dataset identifier PXD067950. Analyzed proteomics data is available in **Source Data 1**. Raw data of all quantifications are available in **Source Data 2**. Any other data reported in this paper will be shared by the lead contact upon reasonable request.

## Supplementary information

### Supplementary videos

Supplementary Video 1. Release of RUSH cargoes in control and under BFA treatment

Supplementary Video 2. Release of RUSH cargoes with P180ΔCC axonal hook

Supplementary Video 3. Release of RUSH cargoes with P180ΔCC axonal hook in desomatized axons

Supplementary Video 4. Axonal ERES are tightly associated with the axonal ER

Supplementary Video 5. Co-movements of SEC22B with ZW10, NBAS and RUSH-SYT1 Golgi-bypassing cargoes

## Source Data

**Source Data 1** (Related to **Fig. 4a** and **Supplementary Fig. 6d and 6e**). Data table containing analyzed proteomics data for SEC13-V5-APEX2 in cortical neurons.

**Source Data 2** (Related to **Figs. 1-7** and **Supplementary Figs. 2, 5, 6, 7, 8 and 9**). Table with the source/raw data for each quantification and uncropped blots presented in the figures and detailed experimental information including N replicates, n number and statistical tests.

