## Supplementary Figures for "The axonal ER couples translation and secretion machineries for local delivery of axonal transmembrane proteins to promote axonal development"

### Supplementary Figure 1

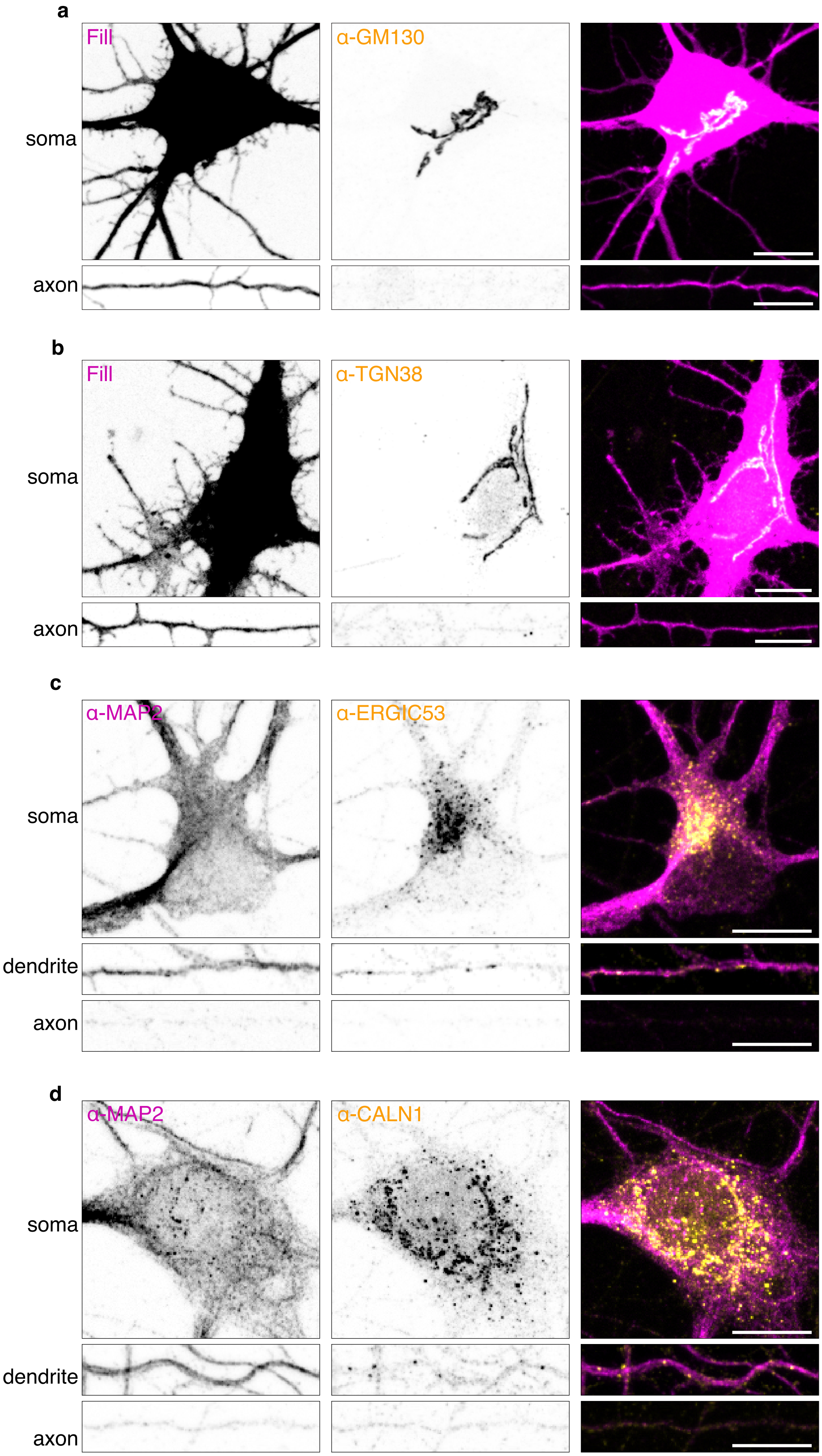

**Supplementary Fig. 1 (Related to Fig. 2): The Golgi and Golgi-related compartments are absent from the axon**

**a, b,** Representative images of neurons DIV9 expressing a fill and stained for endogenous cis-Golgi/Golgi outpost marker GM130 (**a**) or trans-Golgi/Golgi outpost marker TGN38 (**b**) showing their distribution in the somatodendritic domain and absence in the axon. **c,** Representative images of DIV17 human iNeurons stained for the endogenous ERGIC marker ERGIC-53 and the somatodendritic marker MAP2. ERGIC-53 localizes primarily to the soma, a few puncta are observed in the dendrites, but no signal is observed in the axon. **d,** Representative images of DIV10 rat hippocampal neurons stained for the endogenous Golgi satellite marker Calneuron-1(CALN1) and MAP2. CALN1 localizes primarily in the soma and dendrites and is absent from the axons.

Scale bars represent 10 µm in **a, b, c** and **d**.

#### Supplementary Figure 2

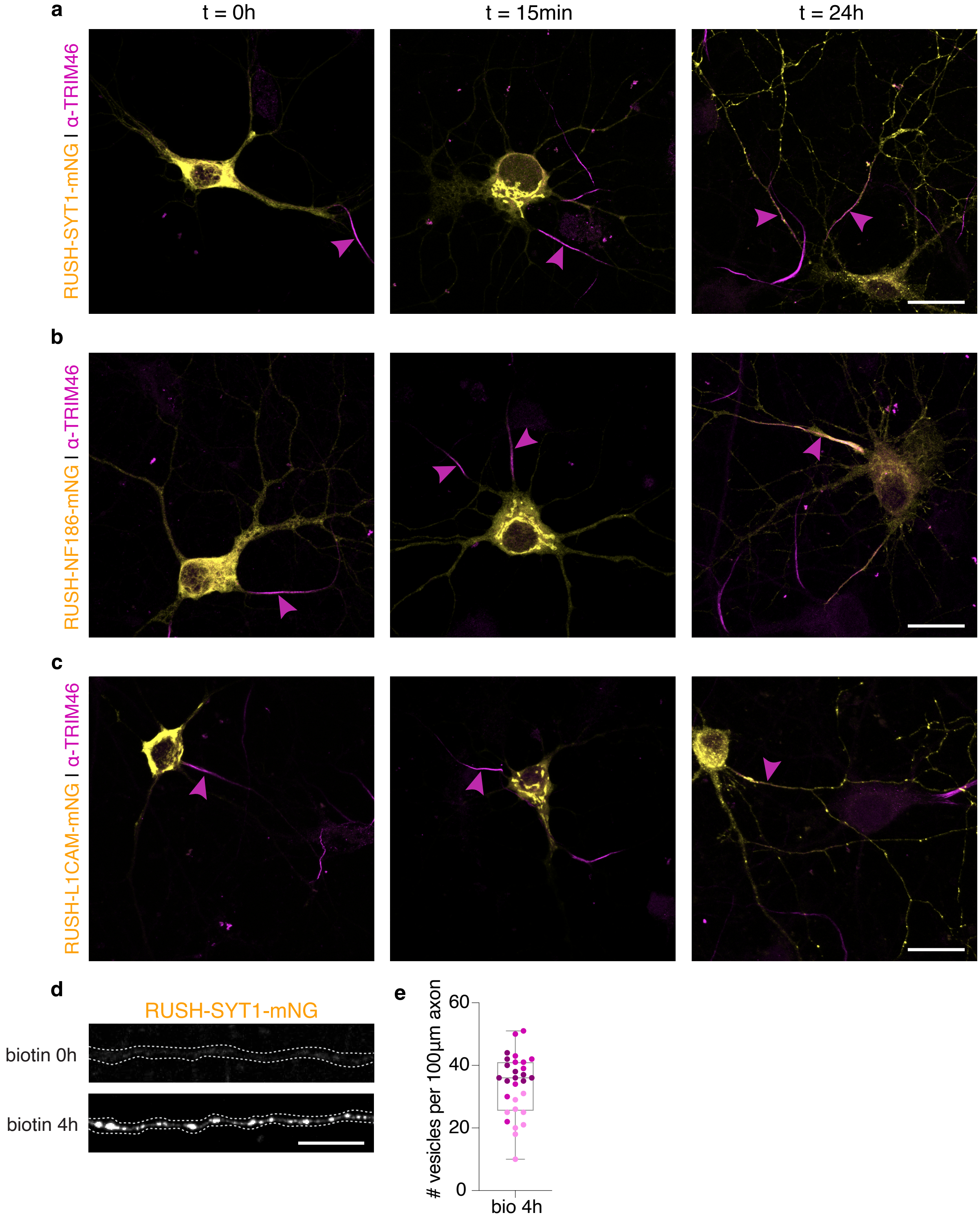

**Supplementary Fig. 2 (Related to Fig. 2): ER retention and trafficking of RUSH cargoes to the PM**

**a-c,** Representative images of neurons DIV10-11 expressing RUSH-SYT1 (**a**), RUSH-NF186 (**b**), or RUSH-L1CAM (**c**) and stained for the endogenous axon initial segment marker TRIM46, showing ER retention at timepoint 0, accumulation at the Golgi after 15 min and localization in target membranes after 24h. SYT1 localized to synaptic vesicles and PM, NF186 enriched at the axon initial segment (AIS) and L1CAM enriched along the axonal PM and in vesicles. **d, e,** Representative images of RUSH-SYT1 cargo distribution in DIV11 axons at timepoint 0 and 4h biotin release (**d**). Quantification of RUSH-SYT1 cargo number per 100 µm axon (**e**) (N=3).

Data are presented as box-and-whisker plot in **e**. Individual data points each represent a neuron, and each color represents an independent experiment. Scale bars represent 20 µm in **a-c** and 10 µm in **d**.

Supplementary Figure 3

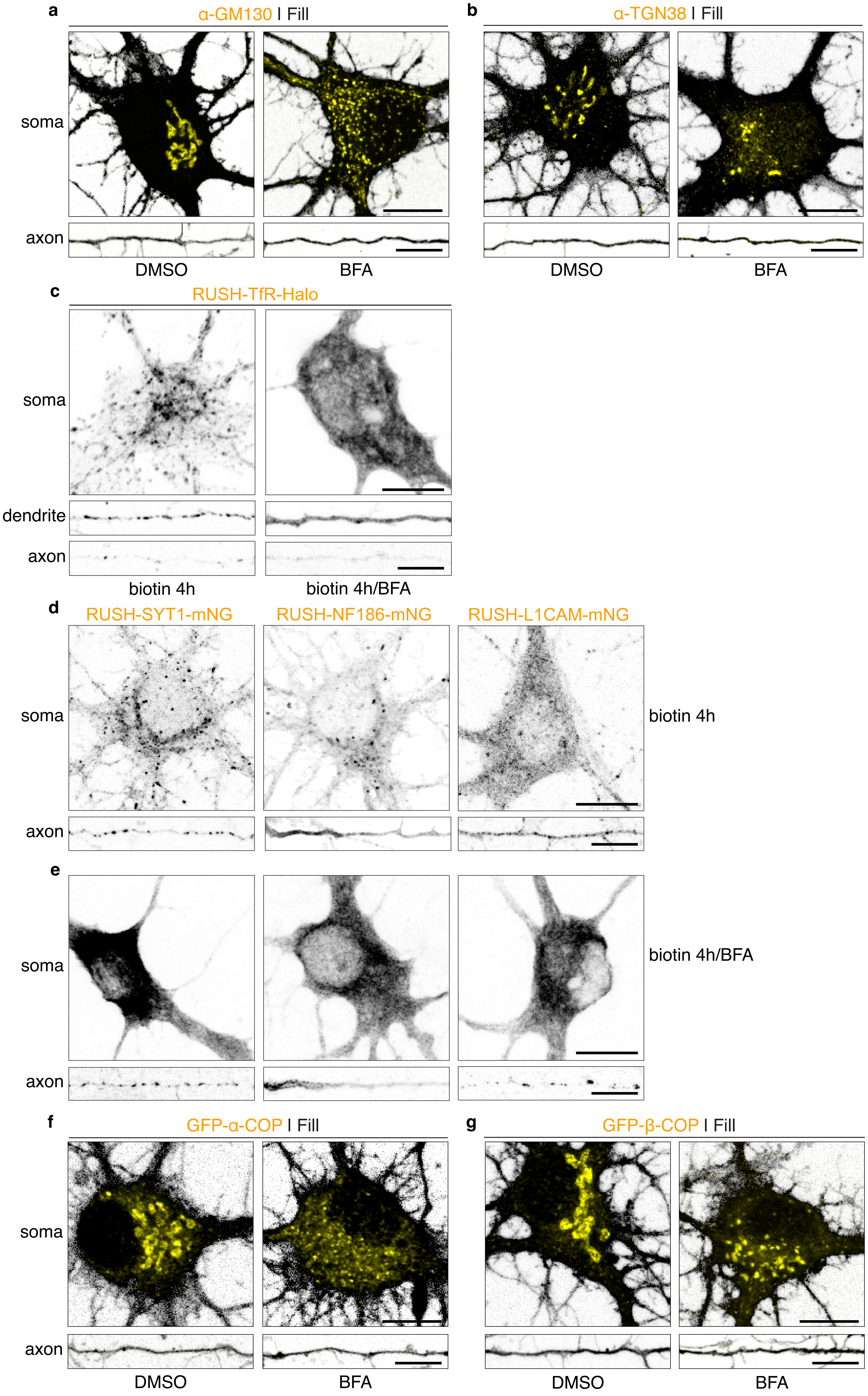

**Supplementary Fig. 3 (Related to Fig. 2): Effect of BFA treatment on Golgi markers and on RUSH cargo release**

**a, b,** Representative images of neurons DIV9-10 transfected with a fill and stained for the cis-Golgi marker GM130 (**a**) or the trans-Golgi marker TGN38 (**a**) in control and after 30 min of BFA treatment. Images show intact Golgi in control condition, and Golgi fragmentation after BFA treatment. In both conditions, Golgi markers were absent in the axon. **c,** Representative images of RUSH-TfR cargo distribution in the soma, dendrite and axon after 4h biotin release in neurons pre-incubated or not with BFA for 30 min. **d, e,** Representative images of RUSH-SYT1, RUSH-NF186 and RUSH-L1CAM cargo distribution in the soma and axon after 4h biotin release (**d**) or following 30 min BFA pre-treatment and 4h release (**e**). **f, g,** Representative images of neurons DIV9-10 co-expressing a fill and COPI components α-COP (**f**) or β-COP (**g**) in control and after 30 min of BFA treatment. Images show their localization to the Golgi in the soma in control, association to Golgi fragments after BFA treatment and their absence in the axon.

Scale bars represent 10 µm in **a-g**.

### Supplementary Figure 4

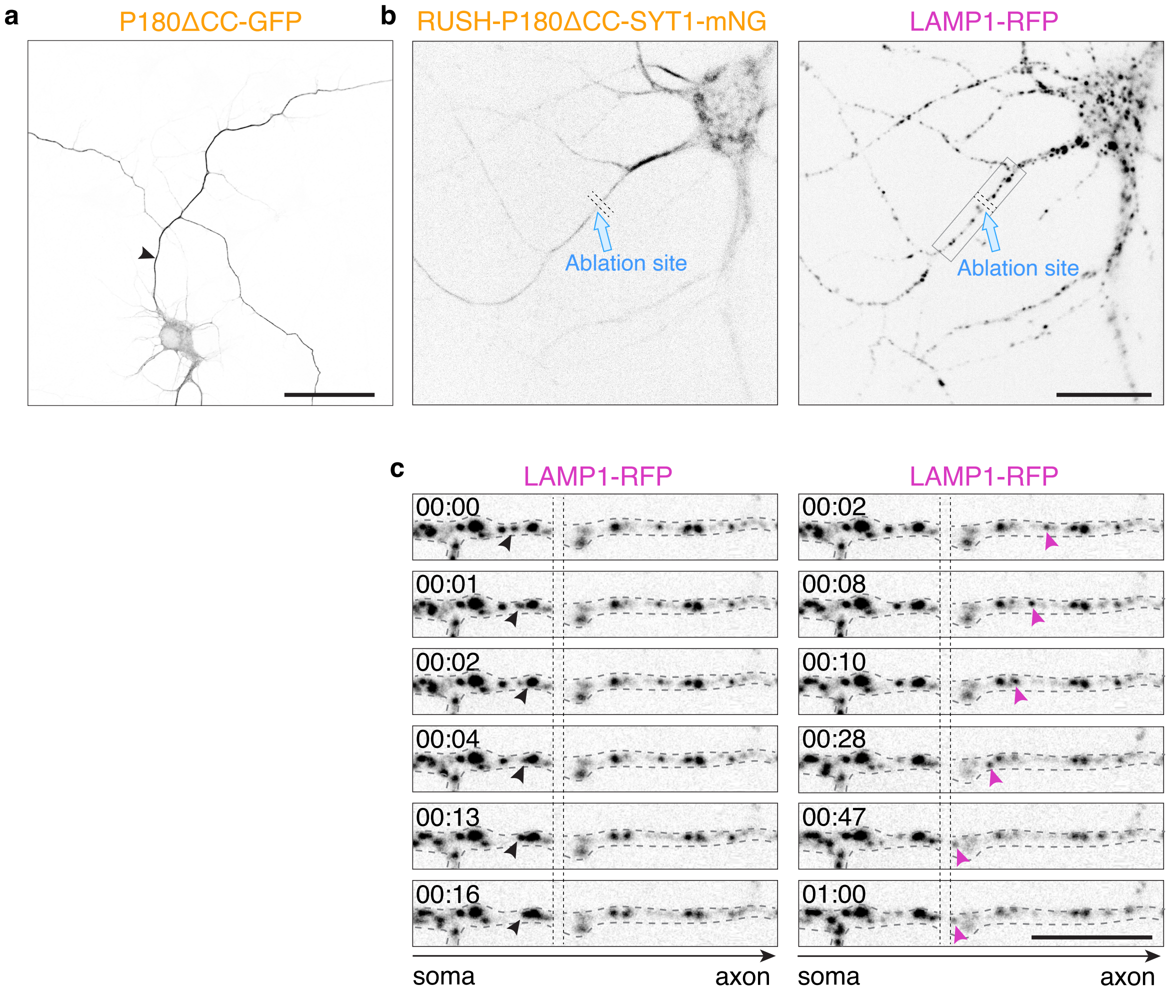

**Supplementary Fig. 4 (Related to Fig. 2): Axonal P180ΔCC localization and validation of soma-axon photoablation**

**a,** Representative image of a neuron DIV6 expressing axonal enriched P180ΔCC construct, arrowhead points to the axon. **b,** Representative images of a neuron transfected with RUSH-P180ΔCC-SYT1 and LAMP1-RFP, which were subjected to photoablation. The blue arrows point to ablation site and the rectangle shows the region of axon where live-cell imaging was acquired. **c,** Time series showing movements of LAMP1 in region described in **b**. Arrowheads point to LAMP1 vesicles not being transported between the somatic and axonal regions. Images were acquired at 1-second intervals for 3 min.

Scale bars represent 50 µm in **a**, 20 µm in **b** and 10 µm in **c**.

### Supplementary Figure 5

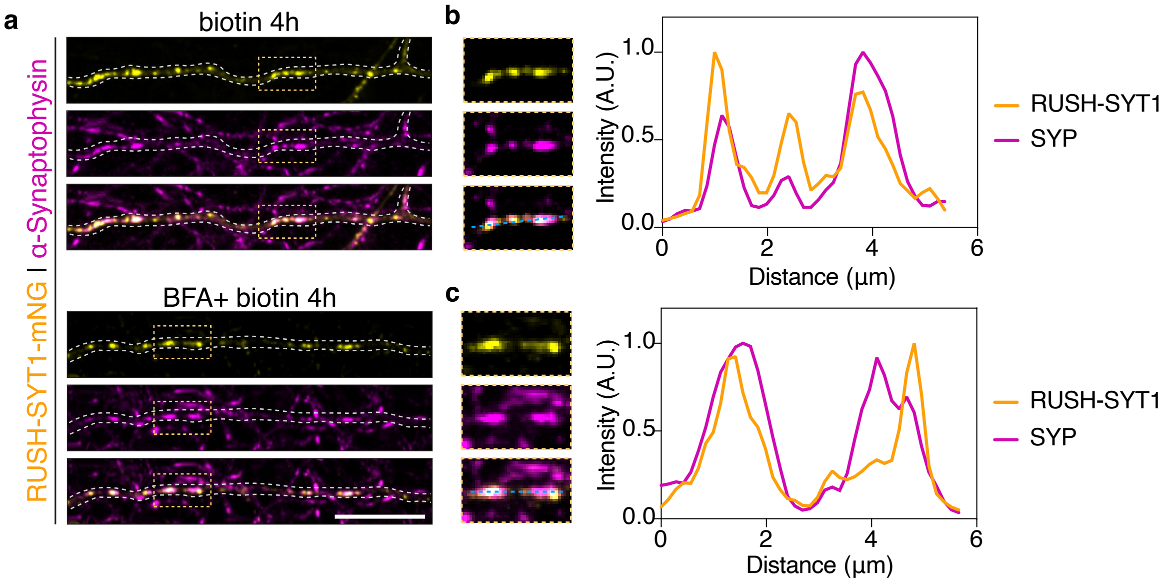

**Supplementary Fig. 5 (Related to Fig. 2): RUSH-SYT1 cargoes reach synaptic vesicles in the presence or absence of BFA**

**a,** Representative images of DIV10 neurons expressing RUSH-SYT1 and stained for endogenous Synaptophysin as a marker for synaptic vesicles. Neurons were treated with either biotin for 4h (left) or pre-treated with BFA for 30 minutes and released with biotin for 4h (right). In both cases, SYT1 cargoes reached synaptic vesicle pools labeled with Synaptophysin. **b, c,** Intensity profile lines from zoomed-in images in **a**, showing colocalization between RUSH-SYT1 and Synaptophysin in control and BFA treated conditions.

Scale bar represents 10 µm.

Supplementary Figure 6

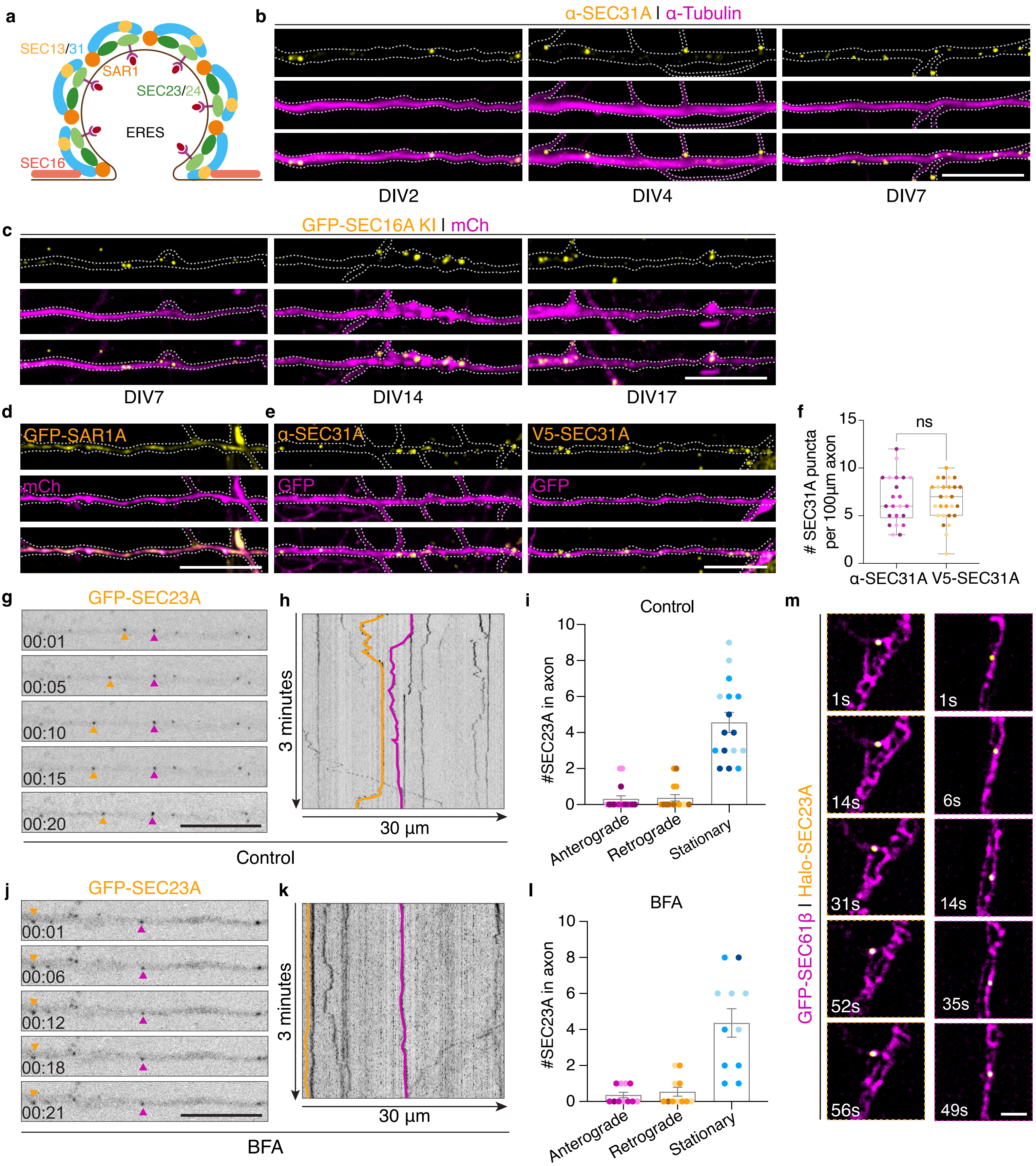

**Supplementary Fig. 6 (Related to Fig. 3)**: **ERES are distributed in axon as stationary complexes across developmental stages**

**a,** Schematic showing essential proteins at the ERES and their organization. **b,** Representative images of axons from rat hippocampal neurons stained for endogenous SEC31 and Tubulin at DIV2, 4 and 7. **c,** Representative images of neurons expressing a fill and a pORANGE knock-in of SEC16A at DIV7, DIV14 and DIV17. **d,** Representative images of a DIV9 neuron expressing a fill and the SAR1A marker. **e, f,** Representative images of neurons DIV9 expressing a fill and stained for endogenous SEC31A or co-transfected with V5-SEC31A (**e**). Quantification of number of endogenous or overexpressed SEC31A puncta per 100 µm axon (**f**) (N=3). **g-l,** Time series of neurons transfected with GFP-SEC23A in control (**g**) or after 30 min BFA treatment (**j**). Images were acquired at 1-second intervals for 3 min. Arrowheads point to examples of SEC23A puncta. Kymographs from axon segments in (**g**) and (**j**) are shown in (**h**) and (**k**), respectively. Quantification of number of anterograde, retrograde or stationary SEC23A puncta in the axon in control (**i**) or after BFA treatment (**l**) (N=3). **m,** Representative time-lapse images of axonal regions showing the tight association between SEC23A and the axonal ER labelled with SEC61β. Images were acquired at 1-second intervals for 80 seconds. See also related Supplementary Video **4**.

Data are presented as box-and-whisker plots in **f** or as mean values ± SEM in **i, l**. Individual data points each represent a neuron, and each color per condition represents an independent experiment. ns = non-significant (p=0.8501) comparing conditions using unpaired t-test in **f**. Scale bars represent 10 µm in **b, c, d, e, g** and **j** and 1 µm in **m**.

Supplementary Figure 7

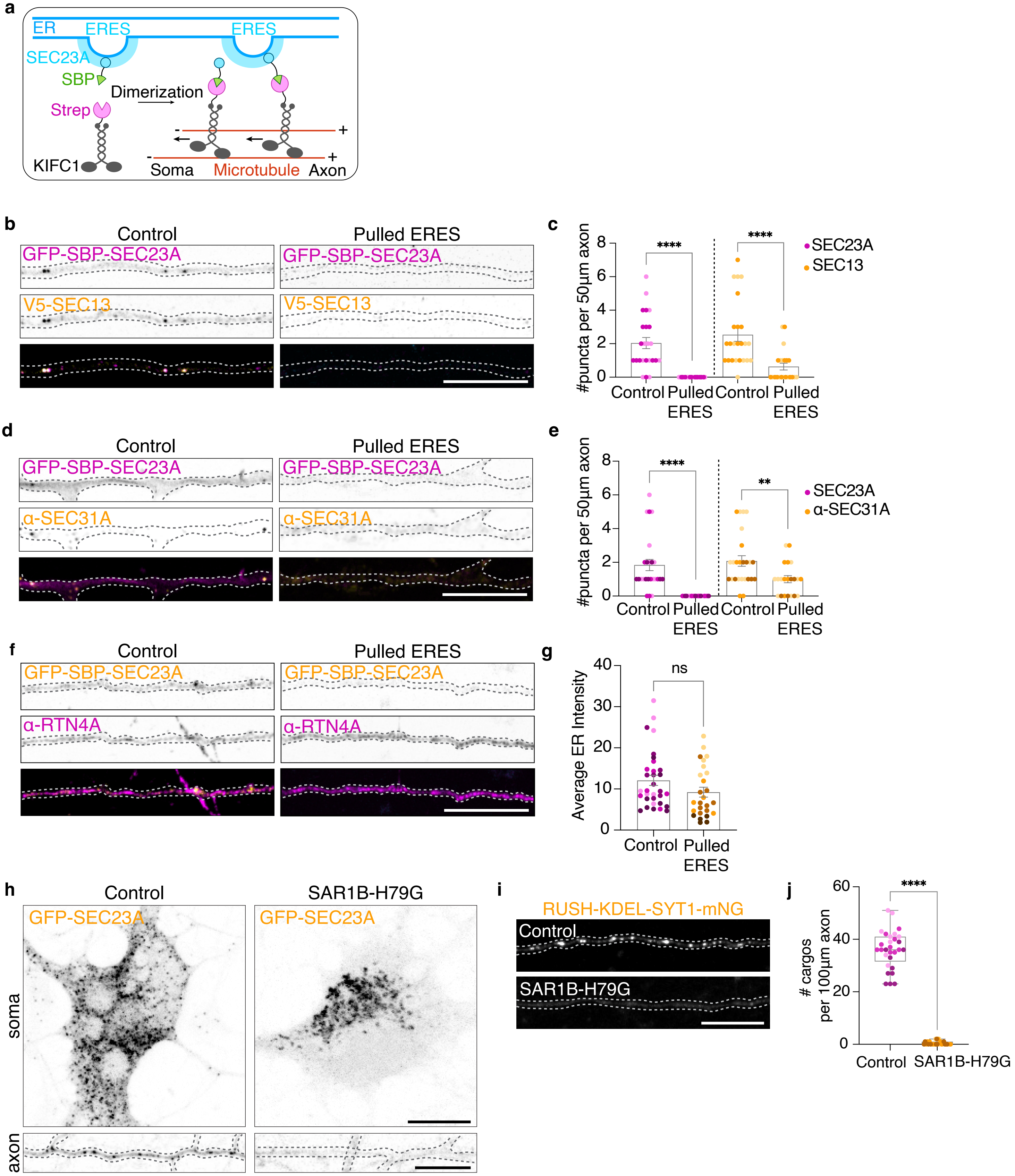

**Supplementary Fig. 7 (Related to Figs. 3, 4 & 6): Tools to assess local function of axonal ERES in axon and effect of ERES function inhibition on cargo exit from the ER**
**a-c,** Schematic showing the Strep-SBP heterodimerization system using Strep-KIFC1 and SBP-SEC23A to relocate axonal ERES to the soma (**a**). Representative images of neurons DIV9 co-transfected with SBP-SEC23A and V5-SEC13 in the absence (control) or presence of Strep-KIFC1 (pulled) (**b**). Quantification of SEC23A and SEC13 puncta in axon (**c**) (N=2). **d,** Representative images of neurons transfected with SBP-SEC23A alone or with Strep-KIFC1 and stained for endogenous SEC31A. **e,** Quantification of SEC23A and endogenous SEC31A puncta in the axon (N=3). **f,** Representative images of neurons transfected with SBP-SEC23A alone or with Strep-KIFC1 and stained for endogenous RTN4 for axonal ER labeling. **g,** Quantification of average ER intensity in the axon (N=4). **h,** Representative images of neurons expressing GFP-SEC23A alone, or with SAR1B-H179G mutant. **i,** Representative images of neurons expressing RUSH-SYT1 alone or with SAR1B-H79G mutant. **j,** Quantification of RUSH-SYT1 cargo number in 100 µm axon in control and in condition where SAR1B-H79G was expressed (N=3).

Data are presented as mean values ± SEM in **c, e, g** or as box-and-whisker plot in **j**. Individual data points each represent a neuron, and each color per condition represents an independent experiment. ns = non-significant ($p=0.0733$) comparing pulled ERES with control condition in **g** using Mann-Whitney test, $^{**}p<0.01$, $^{***}p<0.001$, $^{****}p<0.0001$ comparing conditions to control using Mann-Whitney test in **c, e** (SEC23A) and **j** or unpaired t-test in **e** (α-SEC31A). Scale bars represent 10 µm in **b, d, f, h** and **i**.

### Supplementary Figure 8

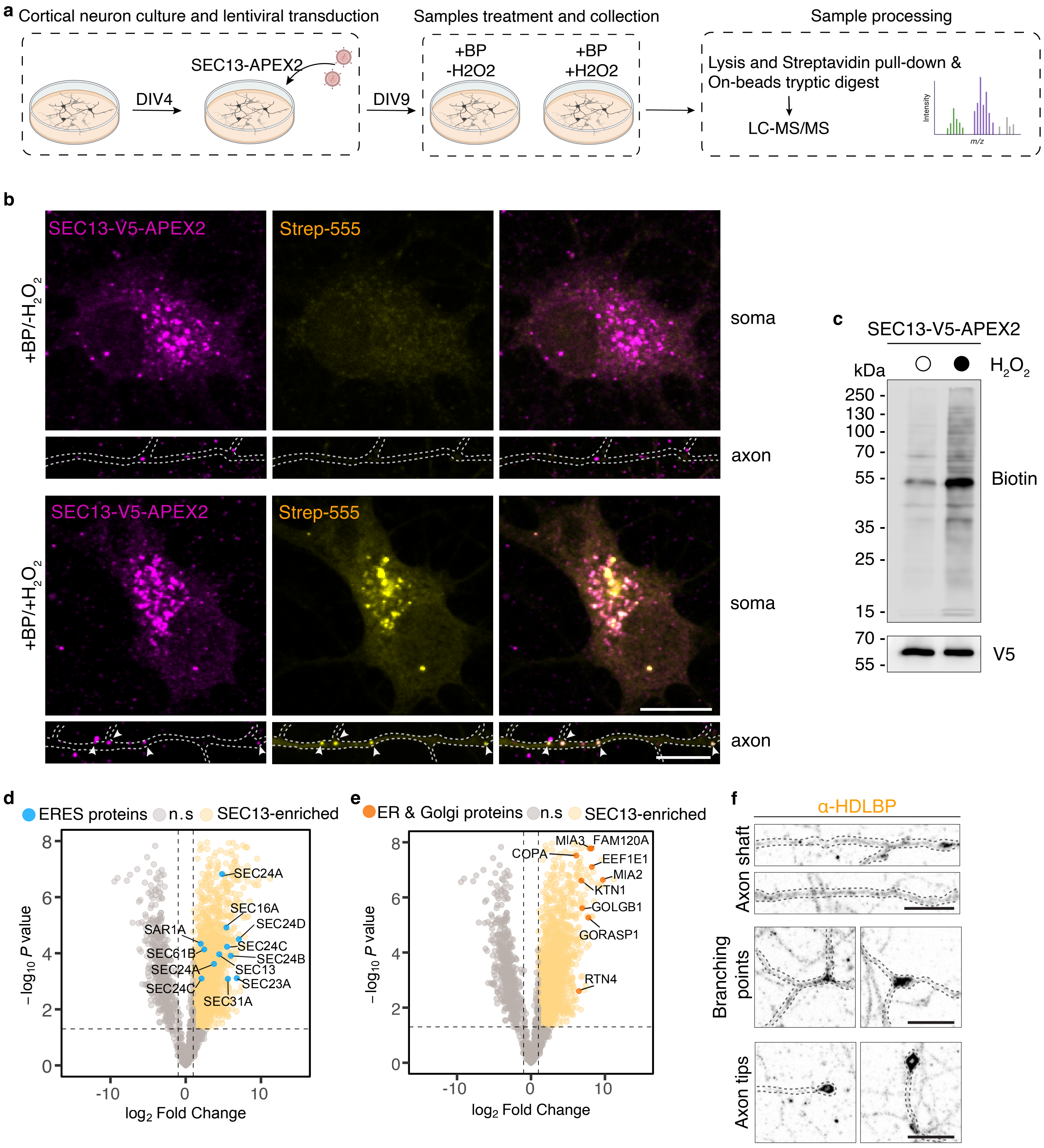

**Supplementary Fig. 8 (Related to Fig. 3). Validation of SEC13-APEX2 tool, and identification of known proteins present in ERES interactome**

**a,** Schematic showing procedure for lentiviral infection of cortical neurons with SEC13-V5-APEX2, sample treatment and mass spectrometry analysis. **b,** Representative images of neurons transduced with SEC13-V5-APEX2, treated with biotin-phenol and with or without $H_2O_2$ as negative control. Expression of SEC13 is visualized with V5 antibody, and biotinylation is detected with conjugated Strep-555. **c,** Immunoblot of SEC13-V5-APEX2 transduced neurons with and without $H_2O_2$. **d, e**, Volcano plots showing the interactome of SEC13 in neurons. Proteins that are enriched near SEC13 are in yellow, other ERES proteins are shown in blue (**d**) and ER and Golgi proteins known to interact with ERES are in orange (**e**). **f,** Distribution of endogenous HDLBP in axon shaft, branching points and axon tips.

Scale bars represent 10 µm in **b** and **f**.

Supplementary Figure 9

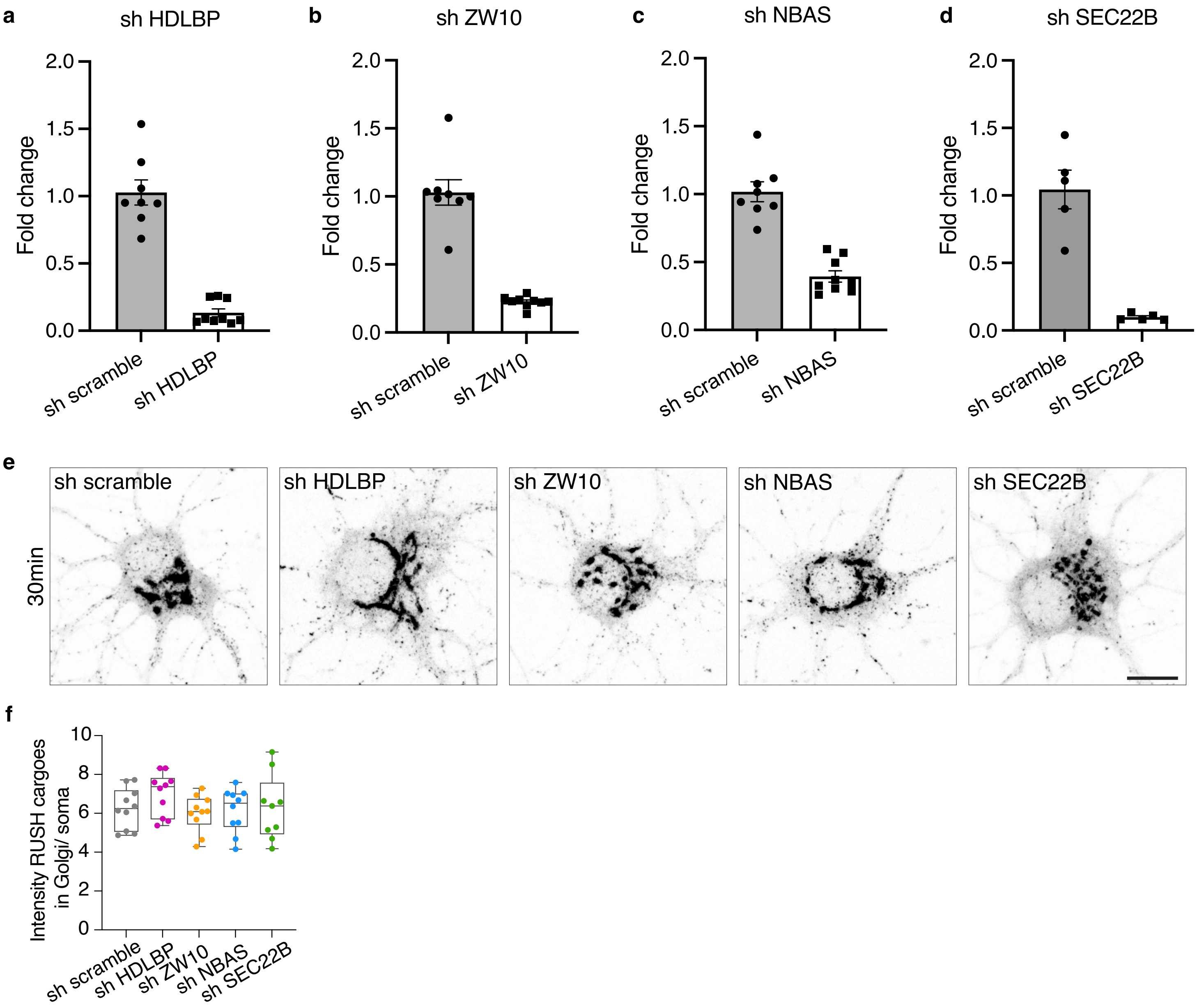

**Supplementary Fig. 9 (Related to Figs. 4 & 5). Validation of shRNA knockdown efficiency for the identified candidate proteins and effect of their knockdown on conventional secretion**

**a-d,** Quantitative PCR fold change of sh HDLBP (**a**), sh ZW10 (**b**), sh NBAS (**c**) and sh SEC22B (**d**) to scramble control. All values were normalized to respective GAPDH internal control from at least 2 experiments with 3 technological replicates per biological sample. **e, f,** Representative images of DIV8 neurons expressing RUSH-SYT1, together with shRNA scramble or shRNA targeting HDLBP, ZW10, NBAS, or SEC22B. Knock-down was performed for 4 days and cargoes were released by addition of biotin for 30 min (**e**). Quantification of intensity of RUSH cargoes in the Golgi, normalized to intensity in the soma (**f**)(N=**1**).

Data are presented as mean values ± SEM in **a, b, c, d** and **h** or as box-and-whisker plots in **f**. Scale bars represent 10 µm in **e**.
